# Structure and mechanism of human dual oxidase 1 complex

**DOI:** 10.1101/2020.09.23.309245

**Authors:** Jing-Xiang Wu, Rui Liu, Kangcheng Song, Lei Chen

## Abstract

Dual oxidases (DUOXs) produce hydrogen peroxide by transferring electrons from intracellular NADPH to extracellular oxygen. They are involved in many crucial biological processes and human diseases, especially in thyroid diseases. DUOXs are protein complexes co-assembled from the catalytic DUOX subunits and the auxiliary DUOXA subunits and their activities are regulated by intracellular calcium concentrations. Here, we report the cryo-EM structures of human DUOX1-DUOXA1 complex in both high-calcium and low-calcium states. These structures reveal the DUOX1 complex is a symmetric 2:2 hetero-tetramer stabilized by extensive inter-subunit interactions. Substrate NADPH and cofactor FAD are sandwiched between transmembrane domain and the cytosolic dehydrogenase domain of DUOX. In the presence of calcium ions, intracellular EF-hand modules enhance the catalytic activity of DUOX by stabilizing the dehydrogenase domain in a position that is optimal for electron transfer.

## Introduction

Reactive oxygen species (ROS) are oxygen-containing chemical species that are highly reactive, such as hydrogen peroxide and superoxide anion ^1^. They participate in many physiological processes and are implicated in several pathological conditions ^1^. ROS can be generated by a class of dedicated enzymes called NADPH oxidase (NOX) in a highly regulated manner. These enzymes are multi-pass transmembrane proteins that catalyze the reduction of extracellular or luminal oxygen by intracellular NADPH to generate superoxide anion or hydrogen peroxide. NOX proteins are involved in many biological processes including host defense, differentiation, development, cell growth and survival, cytoskeletal reorganization and modification of the extracellular matrix ^2^.

Comprising the human NOX protein family are NOX1-5 and DUOX1-2 ^2^. NOX2 protein catalyzes the production of superoxide anion during phagocytosis in neutrophils and is essential for host defense ^3^. DUOX 1-2 proteins are highly expressed in thyroid tissue and they catalyze the production of hydrogen peroxide which is important for the biosynthesis of thyroid hormones^4^. The function of DUOX protein requires physical interactions with an auxiliary protein called dual oxidase maturation factor (DUOXA)^5^. DUOXA promotes the maturation and proper plasma membrane localization of DUOX ^5^. DUOX protein is encoded by two homologous genes in human namely DUOX1 and DUOX2. Similarly, DUOXA protein is encoded by DUOXA1 and DUOXA2. Loss-of-function mutations of DUOX2 or DUOXA2 in human cause congenital hypothyroidism ^6^. Because of the important role of DUOX in thyroid tissue, they are also named thyroid oxidase (ThOX) ^4^.

NOX family proteins share a common catalytic core, formed by a haem-coordinating transmembrane domain (TMD) and a cytosolic dehydrogenase (DH) domain ^7^. DH domain binds intracellular substrate NADPH and cofactor FAD. In addition to the shared TMD-DH catalytic core of NOX, functional DUOX protein has additional large N-terminal extracellular peroxidase homology domain (PHD), a long intracellular Loop 0 containing two EF-hand domains and it requires an auxiliary DUOXA protein for proper function. The activity of DUOX is regulated by intracellular calcium concentration ^4^. Prior to our studies, the structures of NOX family members are only available in the form of isolated domains, including DH domain (PDB ID: 5O0X) ^8^ and TMD (PDB ID: 5O0T) ^8^ of NOX5 from the algea Cylindrospermum stagnale (csNOX5) and a subdomain of human NOX2 DH domain (PDB ID: 3A1F). Despite the functional importance of DUOX and other NOX family members, their structures in the context of full-length functional protein complex are still unknown. Several open questions for DUOX remain elusive: How is the DH domain engaged with TMD to perform the catalytic redox reaction? How does DUOXA protein interact and co-assemble with DUOX? How is the activity of DUOX regulated by intracellular calcium? To answer these fundamental questions, we sought to characterize DUOX-DUOXA protein complex both structurally and functionally.

### Structure determination

To express the human DUOX1 (hDUOX1) protein, we constructed the matured hDUOX1 protein (20-1551) in frame with N-terminal GFP tag guided by rat FSHβ signal peptide for efficient secretion ^9^. The molecular weight of GFP-tagged hDUOX1 is 219 kDa. To monitor the formation of DUOX1-DUOXA1 complex, we fused the human DUOXA1 (hDUOXA1) protein with a C terminal MBP-mScarlet tag to increase its molecular weight to 106 kDa. Fluorescence size exclusion chromatography (FSEC) showed the co-expression of hDUOXA1 effectively shifted the peak of hDUOX1 towards higher molecular weight, suggesting the formation of a stable hDUOX1-hDUOXA1 hetero-complex (Fig.S1a,b). The peak positions indicated hDUOX1 migrated as a monomer, while hDUOX1-hDUOXA1 complex migrated as a hetero-tetramer (Fig.S1b). Moreover, we found the co-expression of DUOX1 and DUOXA1 resulted in cell membranes that showed robust calcium-activated, NADPH-dependent hydrogen peroxide production detected by the Amplex Red assay ^10^ (Fig.1a-c). In the low-calcium condition, DUOX1-DUOXA1 complex showed low basal activity (Fig. 1b,c). Addition of calcium not only reduced the K_m_ but also increased the K_cat_ of DUOX1 complex, leading to the overall enhancement of enzymatic activity (Fig.1b,c).

**Fig. 1.**
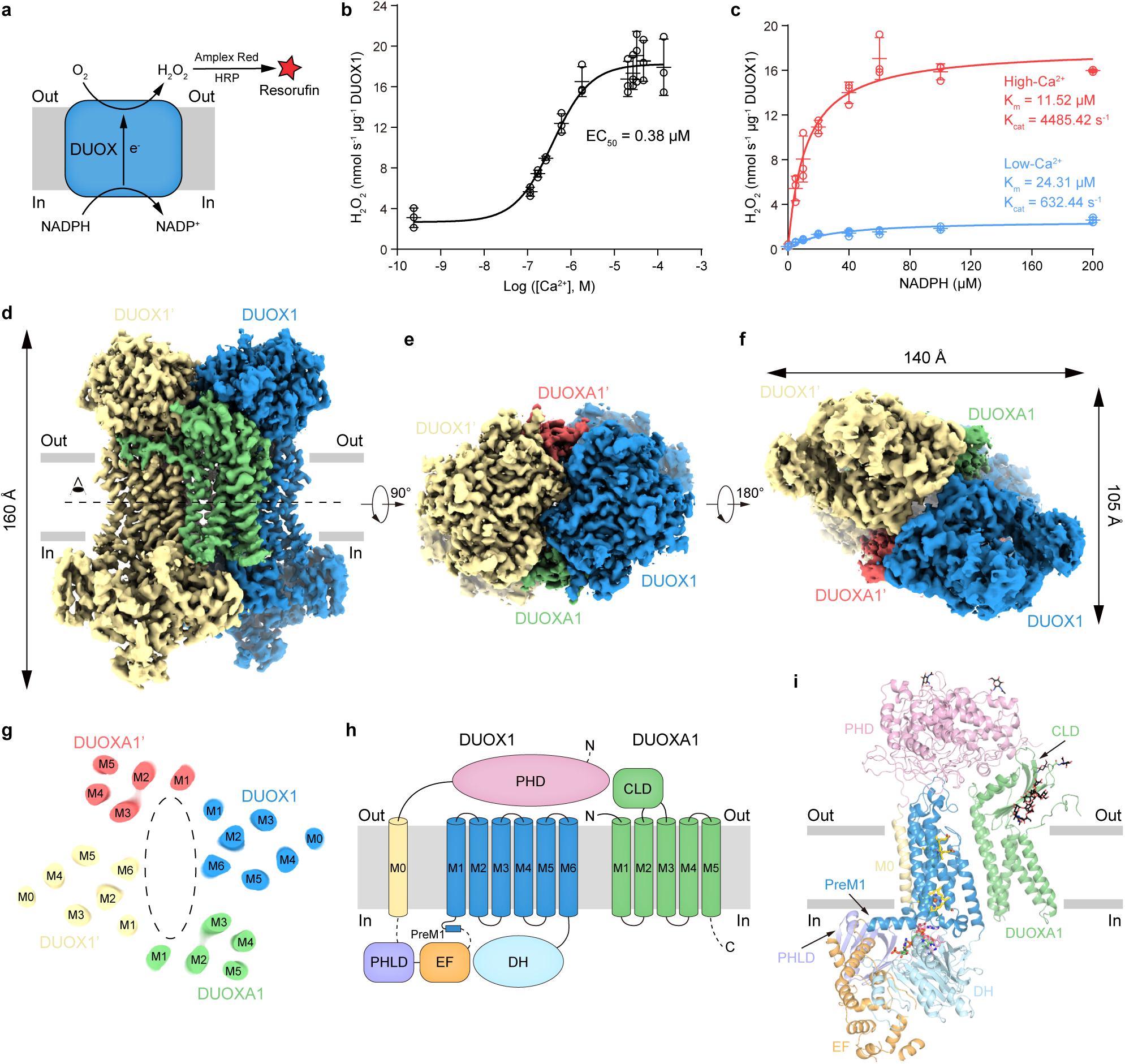
Structure of human DUOX1-DUOXA1 complex in the high-calcium state. **a**, Schematic of the DUOX enzymatic assay. In the presence of H_2_O_2_ (produced by DUOX), horseradish peroxidase (HRP) converts non-fluorescent Amplex Red to fluorescent resorufin, which is measurable and proportional to H_2_O_2_. **b**, Calcium-dependent activation of hDUOX1-hDUOXA1 complex. Data are shown as means ± standard deviations, n = 3 biologically independent samples. **c**, Steady state enzyme activity of hDUOX1-hDUOXA1 complex as the function of NADPH concentration in the presence or absence of calcium. Data were fit to the Michaelis-Menten equation to obtain the K_m_ and K_cat_ value. Data are shown as means ± standard deviations, n = 3 biologically independent samples. **d**, Side view of the cryo-EM map of hDUOX1-hDUOXA1 complex in the high-calcium state. The approximate boundaries of phospholipid bilayer are indicated as gray thick lines. One protomer of hDUOX1 and hDUOXA1 complex is colored as blue and green, the other one is colored as yellow and red, respectively. **e**, A 90° rotated top view compared to **d**. **f**, A 180° rotated bottom view compared to **e**. **g**, Top view of the cross-section of the transmembrane layer at the position indicated as a dashed line in **d**. The large cavity in the transmembrane layer is indicated by dashed oval. For clarity, the cryo-EM map was low-pass filtered to 6 Å. **h**, Topology of hDUOX1 and hDUOXA1 subunits. Transmembrane helices are shown as cylinders, unmodeled disordered regions are shown as dashed lines. The phospholipid bilayer is shown as gray layers. PHD, peroxidase homology domain of hDUOX1; PHLD, pleckstrin homology-like domain of hDUOX1; EF, EF-hand calcium binding module of hDUOX1; DH, dehydrogenase domain of hDUOX1. CLD, claudin-like domain of hDUOXA1. **i**, Structure of one protomer of hDUOX1 and hDUOXA1 complex in cartoon representation. The colors of each individual domain are the same as in **g**. The approximate boundaries of phospholipid bilayer are indicated as gray thick lines. Sugar moieties, haems, FAD and NADPH are shown as black, yellow, pink and green sticks, respectively.

We solubilized and reconstituted hDUOX1-hDUOXA1 complex into peptidisc using NSPr ^11^. Purified peptidisc sample showed a homogenous peak on size exclusion chromatography (Fig.S1c) and two major protein bands on SDS-PAGE, both of which could be trimmed upon PNGase F treatment (Fig.S1d), suggesting both of hDUOX1 and hDUOXA1 were modified by N-linked glycosylation. UV-Vis spectrum showed the peptidisc sample has characteristic Soret band with peak at 415 nm (Fig.S1e), indicating proper Fe(III) haem incorporation. Moreover, the highly purified peptidisc sample recapitulated the calcium-activated NADPH-dependent hydrogen peroxide production observed on membrane (Fig. S1f), confirming that the calcium-dependent activation is a built-in mechanism of hDUOX1-hDUOXA1 protein complex. We prepared cryo-EM grids using the peptidisc sample, either in the presence of 2.5 mM EGTA (low-calcium) or 0.5 mM free calcium (high-calcium). Both samples contained 0.1 mM FAD as cofactor and 0.5 mM NADPH as substrate.

Single particle cryo-EM analysis showed the purified protein was homogeneous and showed two-fold symmetry (Figs.S2-S4). The overall resolution of cryo-EM maps in the low-calcium and high-calcium states reached 2.7 Å and 2.6 Å, respectively (Table S1). The extracellular domains and TMD showed better local resolution than the cytosolic domains, suggesting the higher mobility of the cytosolic domains (Figs.S2g and S4g). To further improve the map quality of cytosolic domains, we exploited symmetry expansion ^12^ and multibody refinement ^13^ by dividing one protomer into the large body (extracellular domain and transmembrane domain) and the small body (the cytosolic domains) (Figs.S2c and S4c). The final resolutions of cytosolic domain reached 3.4 Å and 3.2 Å for the low-calcium and high-calcium states, respectively (Figs.S2-4, Table S1). The high map quality and available homology structures allowed us to build the order regions of the complex which encompassed 88% of DUOX1 and 79% of DUOXA1 (Figs.S5-8, Table S1). In the following text, we will focus on the high-calcium state structure unless noted otherwise, because of its higher resolution.

### The architecture of hDUOX1-hDUOXA1 protein complex

hDUOX1 subunits and hDUOXA1 subunits co-assemble into a 2:2 heterotetrameric protein complex with molecular weight around 457 kDa. The complex encompasses 140 Å ×105 Å ×160 Å 3D space and has an overall two-fold rotational symmetry (Fig.1d-f). Vertically, the complex can be divided into three layers: the extracellular layer, the transmembrane layer and the cytosolic layer (Fig.1d). In the extracellular layer, the two large N-terminal PHD domain of hDUOX1 pack against each other diagonally and are buttressed by the extracellular domain of DUOXA1 from beneath (Fig.1d-f). The transmembrane layer is formed by 24 transmembrane helices and harbors the haem binding sites that provide the electron transfer pathway across the membrane (Fig. 1g). At the center of the transmembrane layer, there is a large cavity without discernable protein densities, which is probably filled by lipids on the cell membrane (Fig. 1g). The cytosolic layer is comprised of the catalytic DH domain and regulatory domains for intracellular calcium sensing (Fig.1f,i).

### Structure of the catalytic hDUOX1 subunit

hDUOX1 is the catalytic subunit of the complex (Fig.2). On the extracellular side of hDUOX1 resides the large N-terminal PHD domain which shares sequence homology with several peroxidases, such as peroxidase A from Dictyostelium discoideum (DdPoxA, PDB ID: 6ERC)^14^ (Fig.S9a). Functional peroxidases utilize histidine-coordinated haem as the cofactor for catalysis. However, key residues for haem binding, such as the haem ligand histidine, are missing in the PHD of hDUOX1. Indeed, we did not observe any haem density in the structure of hDUOX1 PHD, suggesting PHD is probably not enzymatic functional in term of peroxidase activity. Close inspection of the map reveals two putative cation densities in PHD. One cation (cation binding site 1, CBS1) is coordinated by the side chains of D397 and T332, and the main chain carbonyl groups of V399, T332 and R395 (Fig.S9b). The second cation (CBS2) is coordinated by the side chain of D109, D174, S176 and T170 and the carbonyl groups of T170 and W172 (Fig.S9b). We observed strong densities in these two sites in both low-calcium and high-calcium conditions (Fig.S9b), suggesting the bound cations are probably sodium ions which were present in large quantities in our protein sample or calcium ions that bind very tightly. Both CBS1 and CBS2 are evolutionary conserved in DUOX (Fig. S5) and DdPoxA ^14^ (Fig.S9c), indicating their functional importance. Interestingly, we found both CBS1 mutant (D397A+T332A) and CBS2 mutant (D109A+D174A) of DUOX1 failed to co-assemble with DUOXA1 (Fig. S9d). Because CBS1 and CBS2 are away from the subunit interfaces in the DUOX1-DUOXA1 complex, we speculate these mutants probably affect the folding of PHD domain, suggesting the role of CBS1 and CBS2 in protein stability.

**Fig. 2.**
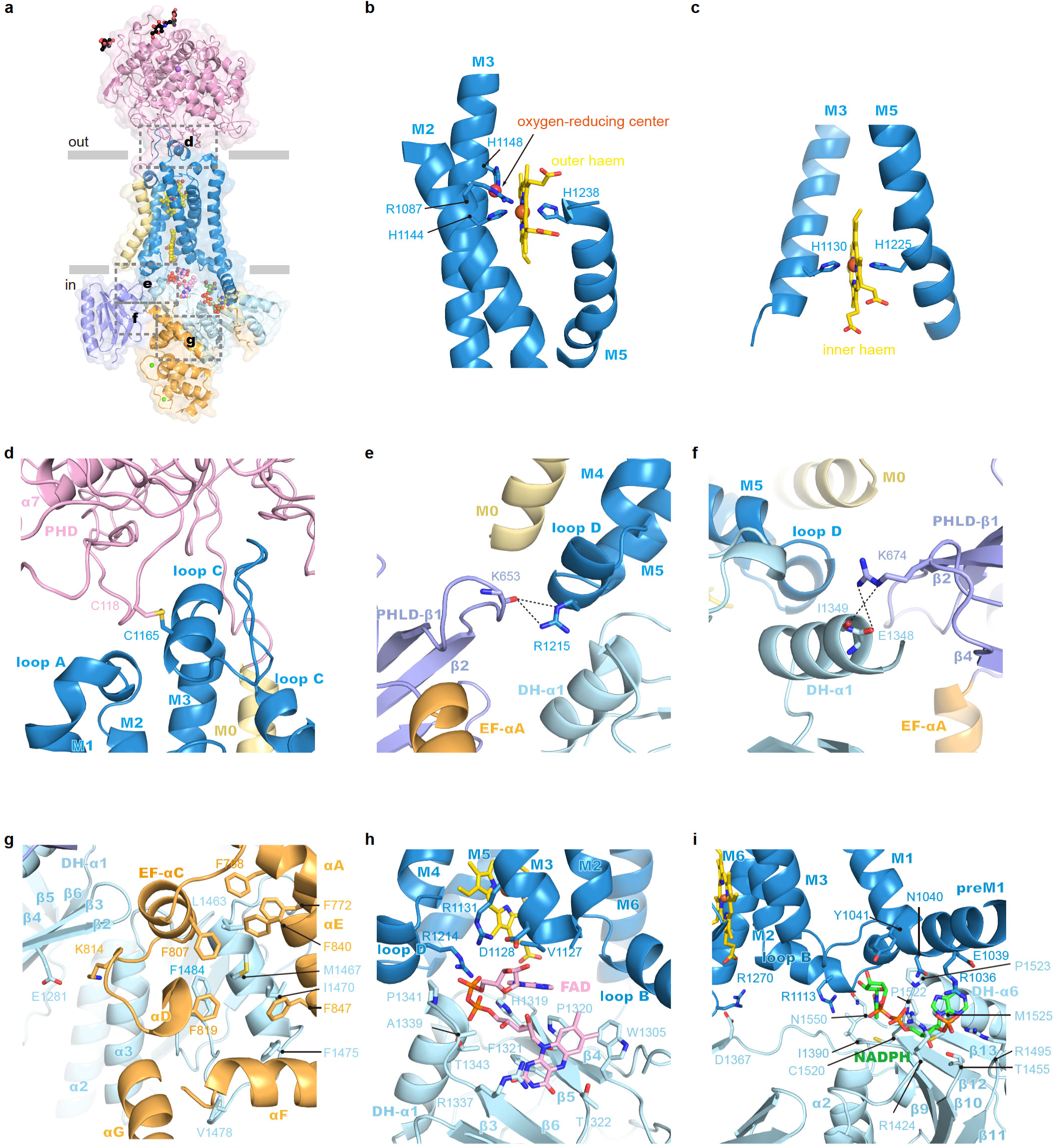
Structure of hDUOX1 subunit. **a**, Side view of hDUOX1 subunit in the high-calcium state, highlighting the key interfaces (boxed by dashed lines). Each domain is colored as in Fig1h. The surface of hDUOX1 is shown in transparency. **b**, The binding site of outer haem in the TMD. Haem is shown as sticks and colored in yellow. Unrelated helices in TMD are omitted for clarity. The putative oxygen-reducing center is indicated by arrow. **c**, The binding site of inner haem in the TMD. **d**, The interface between PHD and TMD boxed in **a**. Disulfide bond between C118-C1165 is shown as golden sticks. **e**, The interface between PHLD and TMD boxed in **a**, the hydrogen bonds are indicated with dashed lines. **f**, The interface between PHLD and DH domain. **g**, The interface between EF module and DH domain. **h**, The FAD binding site located at the interface between TMD and DH domain. Ligands and interacting residues are shown as sticks. **i**, The NADPH binding site located at the interface between TMD and DH domain.

The PHD packs on top of the transmembrane domain of DUOX1 through multiple non-covalent interactions (Fig.2a). Moreover, a disulfide bond between C118 on PHD and C1165 on loop C of TMD further staples the bottom of PHD onto the top of TMD (Fig.2a,d). In the TMD, hDUOX1 has an extra bent M0 helix at the periphery of M3 and M4, together with the canonical 6 TM helices of NOX protein family. M1-M6 of hDUOX1 form two haem binding sites within the transmembrane domain. H1144 on M3 and H1238 on M5 coordinate the outer haem (Fig.2b). H1130 on M3 and H1225 on M5 coordinate the inner haem (Fig.2c). These four histidines are absolutely conserved in NOX family proteins (Fig.S6). We observed a spherical density surrounded by the invariant R1087 on M2, H1148 on M3 and outer haem-coordinating residue H1144 (Fig.2b and S3c). Previous studies showed mutations of the csNOX5 residues corresponding to R1087 and H1148 of hDUOX1 affected the re-oxidation of dithionite-reduced TMD by oxygen and this site was proposed to be the oxygen substrate binding site, namely oxygen-reducing center (Fig.S10a,b) ^8^. Our structure observations in hDUOX1 support the hypothesis.

Preceding the M1 helix of DUOX1 TMD, an amphipathic preM1 helix floats on the inner leaflet of plasma membrane (Fig.1h). This helix was previously observed in csNOX5 ^8^ and is probably a shared feature of NOX family proteins. Between M0 and pre-M1 is a long cytosolic fragment loop 0. Cryo-EM maps reveal that the N terminal of loop 0 is a domain rich of β sheets (Fig.S3g and S10c). Structural search using DALI server ^15^ identified the β sheets-rich domain is a crypto pleckstrin homology-like domain (PHLD) that shares little sequence homology but high structural similarity to the pleckstrin homolog (PH) domain proteins (Fig.S10c)^16^.

Following the PHLD, two EF-hand type calcium binding domains (EF1 and EF2) form a compact helical module that is connected to the PHLD through αC (Fig.S6). Residues predicted to be responsible for calcium binding in EF1 and EF2 are evolutionary conserved in DUOX family proteins (Fig.S6). Although we did not observe the strong densities for small calcium ions due to poor local resolution (Figs.S2-3), the structure of EF-hand module closely resembles the small subunit of calcium-dependent protein phosphatase calcineurin in calcium-bound state (PDB ID: 4IL1) ^17^ (Fig.S10d), suggesting both EF1 and EF2 are loaded with calcium in the high-calcium state. Based on the homology structure (4IL1), side chains of D828, D830, N832 and E839 and the main chain carbonyl group of Y834 chelate one calcium ion in EF1 (Fig.S10e) and side chains of D864, D866, N868, E875 and the main chain carbonyl group of L870 chelate another calcium ion in EF2 (Fig.S10f). It is reported that mutations of any of these calcium binding sites abolished calcium activation ^18^ and E879K mutation in hDUOX2 (E875 in hDUOX1) lead to congenital hypothyroidism ^19^, emphasizing their importance in calcium activation.

The C-terminal catalytic DH domain is connected to M6 of DUOX1 TMD via a short linker (Fig.1h,i). DH domain of hDUOX1 has a canonical dehydrogenase fold and its structure is similar to csNOX5 ^8^ (Figs.1i and S7). We observed strong densities for both FAD cofactor and NADPH substrate and their binding sites were contributed from not only DH domain but also TMD (Fig.2h,i), as described later.

### Inter-domain interactions in the high-calcium state stabilize the electron transfer pathway

In the high-calcium state, individual domains of DUOX1 in the cytosolic layer are stabilized by multiple inter-domain interactions. The PHLD interacts with adjacent TMD and DH domains (Fig.2e). The main chain carbonyl group of K653 on PHLD makes hydrogen bond with R1215 on loop D of TMD (Fig.2e). Side chain of R674 of PHLD interacts with the main chain carbonyl group of E1348 and I1349 on α1 of DH domain (Fig.2f). The EF1-EF2 module in the high-calcium state shapes a crevice that embraces α4 and post α4 loop of DH domain (Fig.2g and Fig.S10g). The interactions between EF module and DH are mainly hydrophobic and involve F768, F772, F807, F819, F840 and F847 of EF module, L1463, M1467, I1470, F1475, V1478 and F1484 of DH domain (Fig.2g). In addition, K814 of EF module makes electrostatic interaction with E1281 on β2 of DH (Fig.2g). The interactions between EF module and DH domain of hDUOX1 mimic the interactions between calcineurin subunit B and A in the calcium-bound state (PDB ID: 4IL1)^17^ (Fig.S10g,h).

The linker between EF module and preM1 helices binds in a groove on the surface of DH domain (Fig.1i). DH docks onto the bottom of TMD via polar interaction between R1270 on M6 and D1367 on β7, and between R1113 on loop B of TMD and N1550 of DH (Fig.2h,i). It is reported that R1111Q mutation in hDUOX2 (R1113 in hDUOX1) was identified in congenital hypothyroidism patients ^19^, highlighting the importance of this inter-domain interaction. Moreover, both the FAD cofactor and NADPH substrate bind at the interface between DH and TMD. R1214 and R1131 in TMD form electrostatic interaction with phosphate of FAD. D1128 makes hydrogen bonding with ribose of FAD (Fig.2h). E1039 and N1040 in TMD make hydrogen bonding with adenosine ring of NADPH, and R1036 make cation-π interaction with both adenosine ring and electrostatic interaction with phosphate group of NADPH (Fig.2i). Notably, R1495, R1424 and R1036 all participate in electrostatic interactions with the phosphate group of NADPH ribose, providing structural mechanism to distinguish NADPH from NADH (Fig.2i). Through structural comparison, we found the NADPH binding site in csNOX5 structure was blocked by the artificially engineered C-terminal insertion which was introduced into previous crystallization construct ^8^ (Fig.S10i). Moreover, the adenosine group of FAD has a 180° flip compared with structure of isolated DH of csNOX5 (Fig.S10j), suggesting the TMD has strong influence on the binding and conformation of the FAD molecule. Taken together, the binding of FAD and NADPH onto hDUOX1 would bridge and stabilize the docking of DH onto TMD.

The aforementioned inter-domain interactions involve a complex nexus among PHLD, EF-hand module, DH domain and TMD, and stabilize DUOX1 in a catalytic competent conformation for efficient electron transfer. In this conformation, the measured edge-to-edge distances between NADPH and FAD, between FAD and inner haem, and between inner haem and outer haem are 8.2 Å, 3.9 Å and 6.7 Å, respectively (Fig.3a). It is possible that there are additional protein residues on DUOX1 that also participate in the electron rely process. At the end of electron transfer chain, the initial product of oxygen reducing reaction is superoxide anion. We probed the possible pathways for oxygen entrance and for superoxide anion exit with CAVER ^20^, using oxygen-reducing center as the starting point. We located four possible tunnels: tunnel A is formed by M1, M2, M5 and M6 and is capped by loop E on top (Fig.3b); tunnel B is surrounded by M2, loop A, loop C and loop E (Fig.3c); tunnel C is embraced by M3, M4 and loop C (Fig.3d); tunnel D is enclosed by M3, M4, loop C and M0 (Fig.3e). The bottleneck radii of these tunnels are around 1Å (Fig.3f), which may allow the permeation of small oxygen substrate under dynamic motion of DUOX1 protein. Further analysis showed tunnel B-D are all surrounded by hydrophobic residues (Fig.3c-e), which are unfavorable for superoxide anion permeation. In contrast, tunnel A is gated by hydrophilic R1087 on M2, R1062 on M1, R1248 and Q1245 on loop E (Fig.3b). We speculate the highly positively charged constriction of tunnel A would strongly attract the negatively charged superoxide anions, and this might be essential for the dismutation reaction between two superoxide anions to generate uncharged hydrogen peroxide for diffusion. Therefore, manipulations that may alter the constrictions of tunnel A-D would affect superoxide anion intermediate leakage. Indeed, it is reported that mutations on DUOX1 loop A or on DUOXA1 NTP which interacts with and stabilizes loop A would change the ratio of superoxide anion and hydrogen peroxide produced, probably by affecting the leakage of superoxide anions through these tunnels ^21–23^.

**Fig. 3.**
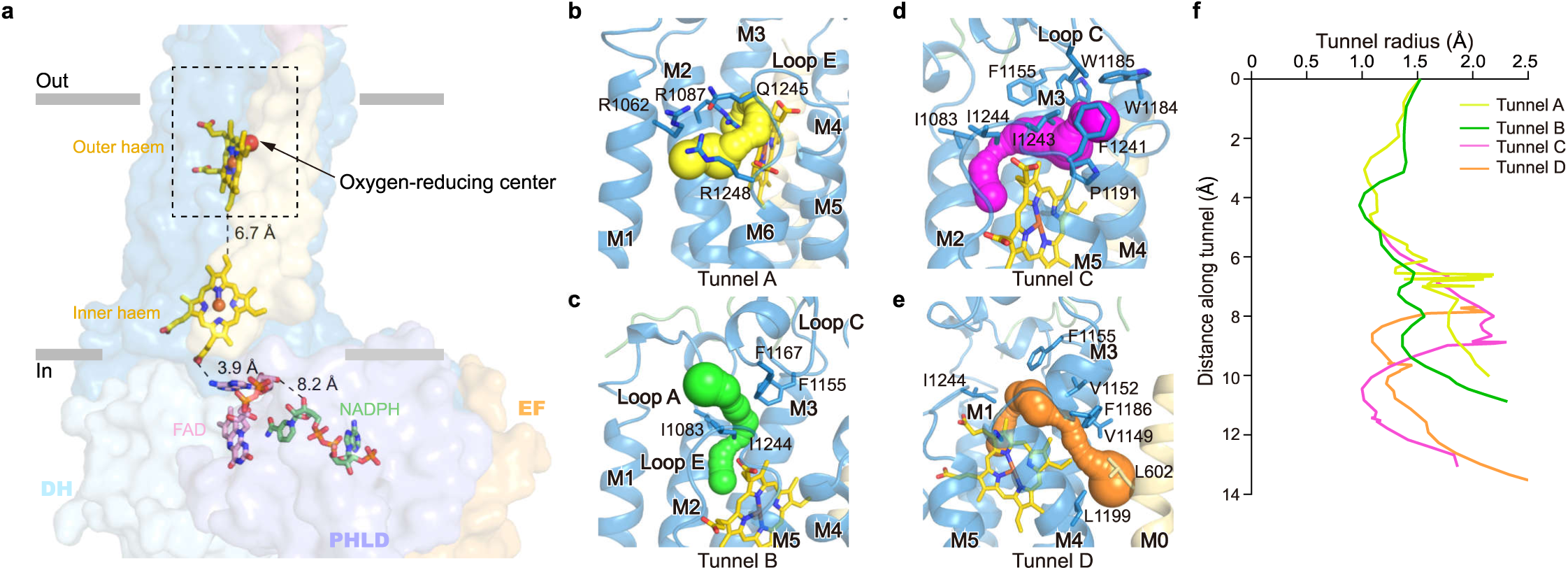
Electron transfer pathway in hDUOX1 subunit in the high-calcium state. **a,** The edge-to-edge distances between NADPH and FAD, FAD and inner haem, and two haems are shown beside dashes. The ligands are shown as sticks, each domain of hDUOX are shown in surface, and colored the same as Fig. 1h. Only one hDUOX subunit is shown for clarity. The putative oxygen-reducing center is boxed by dashed lines. **b-e**, The close-up view of the putative oxygen-reducing center. Four predicted tunnels for oxygen substrate entrance and product exit are shown as surface in yellow, green, magenta and orange, respectively. Residues surrounding the tunnels are shown as sticks. **f**, Calculated radii of tunnels shown in **b-e**. The putative oxygen-reducing center is used as the starting point for calculation.

### Structure of hDUOXA1 and mechanism of complex assembly

DUOXA protein is an essential auxiliary subunit for DUOX enzyme ^5^ (Fig.S11a) and it has an extracellular N-terminus that is important for hydrogen peroxide generation ^22, 23^. We observed the N-terminus peptide (NTP) of hDUOXA1 extends and packs onto the PHD-TMD junction of the distal hDUOX1 subunit (Fig.4a-g). Side chains of F8, F10 and Y11 of NTP insert into the hydrophobic groove formed by loop C, loop A and PHD of hDUOX1 (Fig.4c). In addition, K15 of DUOXA1 NTP makes electrostatic interactions with D1077 of DUOX1 (Fig.4d). This agrees with previous data showing DUOXA1 NTP interacts with DUOX1 loop A ^21^. hDUOXA1 has five transmembrane helices. Lower part of TM1 interacts with preM1 and M1 of hDUOX1 (Fig.S11a). The remaining four helices and associated extracellular loops share structural similarity with claudin superfamily members, such as claudin-9 (PDB ID:6OV2)^24^ (Fig.S11). The extracellular loop between TM2 and TM3 folds into a compact claudin-like domain (CLD) composed of four β strands and two α helices (Fig.S8 and S11). CLD forms extensive interactions with both distal and proximal DUOX1 subunits (Fig.4a,b), emphasizing its important role in the complex assembly. This agrees with previous studies showing that splicing variants at TM2-TM3 loop have distinct behavior in supporting the activity of DUOX1 ^22^. Moreover, we found an ordered N-linked glycosylation decoration on N109 of hDUOXA1 and its branched sugar moieties make extensive polar interactions with both DUOXA1 and DUOX1 subunits (Fig.4a,b). The PHD of two DUOX1 subunits also interact with each other (Fig.4a,b). Close to the dyad axis, R50 and R507 on one PHD make polar interactions with E41 and F313 on the opposite PHD (Fig.4e). We further analyzed the effects of interface mutations on the tetramer assembly and found mutations of R50E, R507E and R507A all severally affect tetrameric peak formation on FSEC (Fig.4f). These structural information and biochemical data revealed the detailed inter-subunit interactions that dictate the hetero-tetramer assembly.

**Fig. 4.**
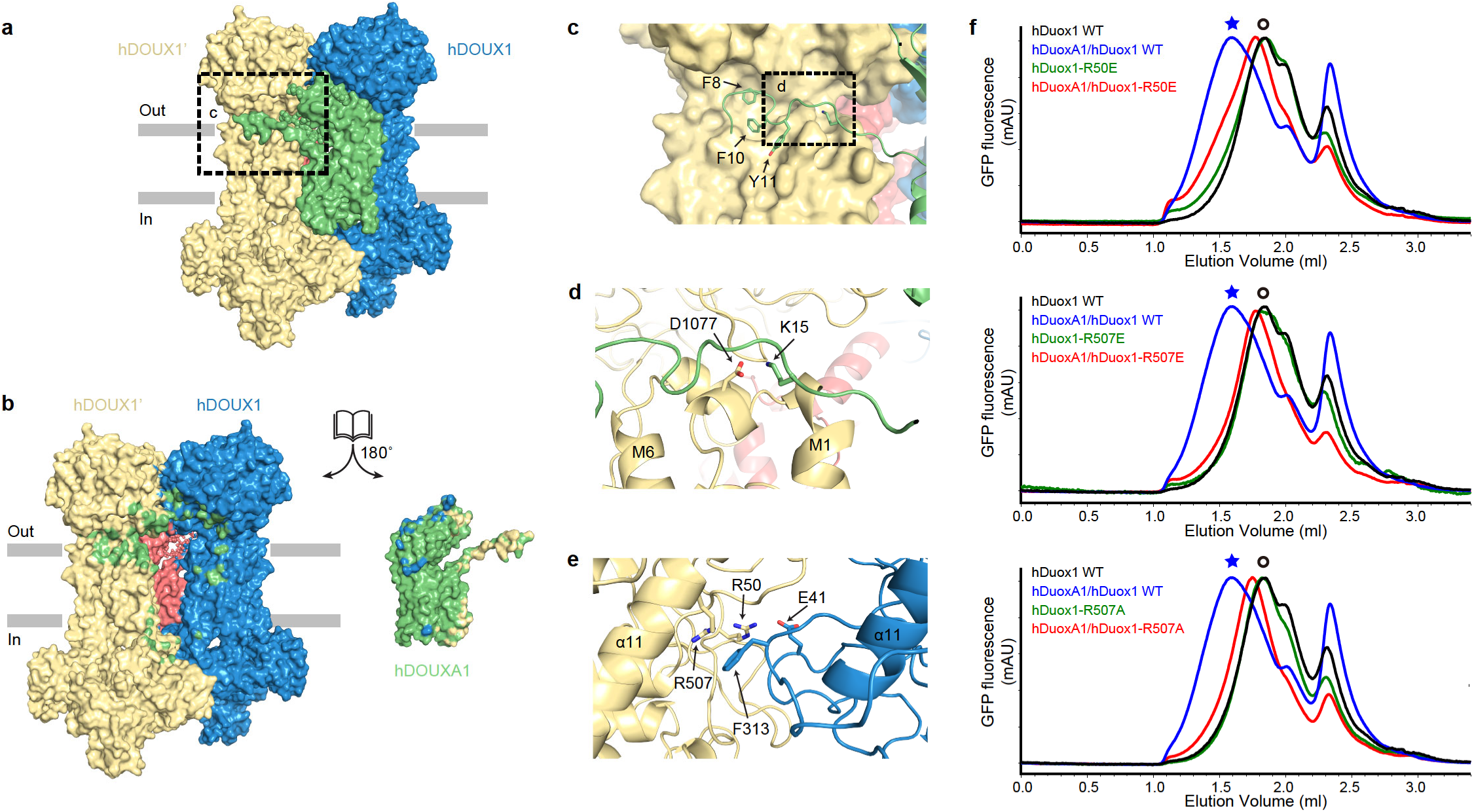
Mechanism of hDUOX1-hDUOXA1 tetramer assembly. **a**, The side view of hDUOX1-hDUOXA1 protein complex shown in surface representation and colored the same as in Fig.1d. **b**, The open-book view of the inter-subunit interfaces. Residues of hDUOX1 subunits that interact with hDUOXA1 subunit are colored in green. Residues of hDUOXA1 subunit that interact with hDUOX1 subunits are colored in yellow and blue. **c,** The close-up view of the interactions between NTP of hDUOXA1 and hDUOX1 boxed in **a**. **d,** The close-up view of additional interactions between NTP of hDUOXA1 and hDUOX1 boxed in **c**. **e**, The top view of interactions between PHD of two opposing hDUOX1 subunits. **f**, Representative FSEC traces of hDOUX1 R50E, R507E and R507A mutants are compared to that of wild-type (WT) hDOUX1. The peak position of the hDOUX1 peak is denoted by the hollow circles. Asterisks denote the peak position of hDUOX1-hDUOXA1 protein complex.

### Activation mechanism of DUOX1 complex by calcium

The consensus map in the low-calcium state showed the cytosolic layer had poor local resolution which was improved by multibody refinement ^13^ (Fig.S4). Further molecular flexibility analysis ^13^ showed the cytosolic domains (small body) in the low-calcium state were sampling a broad range of orientations relative to the transmembrane domain, evidenced by the plateau-shaped distribution on the histogram of the major eigenvector (Fig. S4f). This is in great contrast to the normal distribution in the high-calcium state (Fig. S2f), suggesting the cytosolic layer in the low-calcium state is more flexible. We compared the structures in the low-calcium state and high-calcium state and found structural changes in the extracellular layer and transmembrane layer are small (Fig.5a). However, there are large conformational changes of the regulatory PHLD and EF-hand module in the cytosolic layer (Fig.5a-c, Movie S1). In the absence of calcium, the EF module switches from an extended shape into a more contracted shape (Fig.5d-e), which reconfigures the interface between EF module and α4 of DH domain, resulting in a loosely packed structure (Fig.5f). In the low-calcium state, EF2 moves away from DH domain. The Cα atom of A894 on αJ of EF2 has 40 Å displacement (Fig. 5b). PHLD rotates away from the TMD and DH domain and αA of PHLD has 17.2° outward rotation (Fig.5c). As a result, several inter-domain interactions observed in the high-calcium state were disrupted and therefore the docking of DH domain onto TMD is weakened by these structural changes, leading to a higher mobility of DH domain (Fig.S4g). We propose the increased mobility of DH domain negatively correlates with the electron transfer efficiency and thus the catalytic activity of DUOX. In addition, because TMD also contributes to FAD and NADPH binding, the increased mobility of DH domain would result in reduced affinity of NADPH as well. This is in agreement with the markedly reduced K_cat_ and moderately increased K_m_ in the low-calcium state as we observed (Fig.1c).

**Fig. 5.**
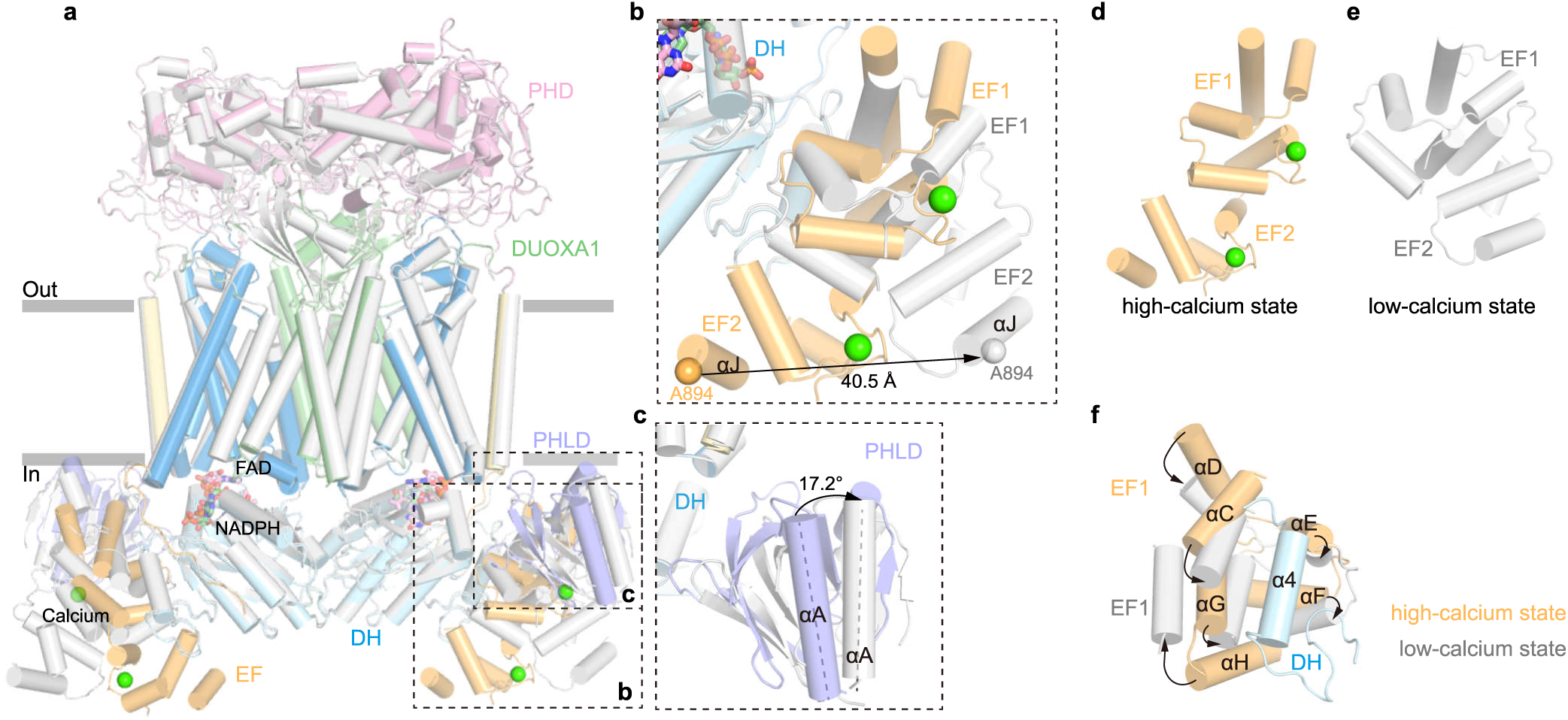
Conformational change of hDUOX1 complex during calcium activation. **a**, Structural comparison of hDUOX-hDUOXA1 complex between the high-calcium state (colored) and the low-calcium states (gray). Protein is shown as cartoon. Regions with large conformational changes are boxed by dashed lines. **b**, Close-up view of the conformational changes of EF-hand module. Cα atom of A894 on αJ helix is used as marker to measure the movement of EF2. **c**, Close-up view of the conformational change of PHLD. The angle between αA helices in the high-calcium and low-calcium states was measured. **d-e**, Conformational differences of EF-hand module between the high-calcium state and the low-calcium state. **f**, Reconfiguration of the interface between EF-hand module and α4 helix of DH domain. Arrows denote movements from high calcium state into the low-calcium state.

## Conclusions

In this study, we provided the structures of hDUOX1-hDUOXA1 as a hetero-tetrameric protein complex in both high-calcium state and low-calcium state. The structure of hDUOX1 complex in the high-calcium state reveals multiple inter-domain interactions that optimally orientate DH domain and TMD for efficient electron transfer and thus redox reaction. Removal of calcium ions results in the reconfiguration of cytosolic inter-domain interactions which in turn mobilizes the DH domain, and lowers the electron transfer efficiency (Fig.6). These structures provide mechanistic insights into the structure and mechanism of DUOX and other NOX enzymes.

**Fig.6.**
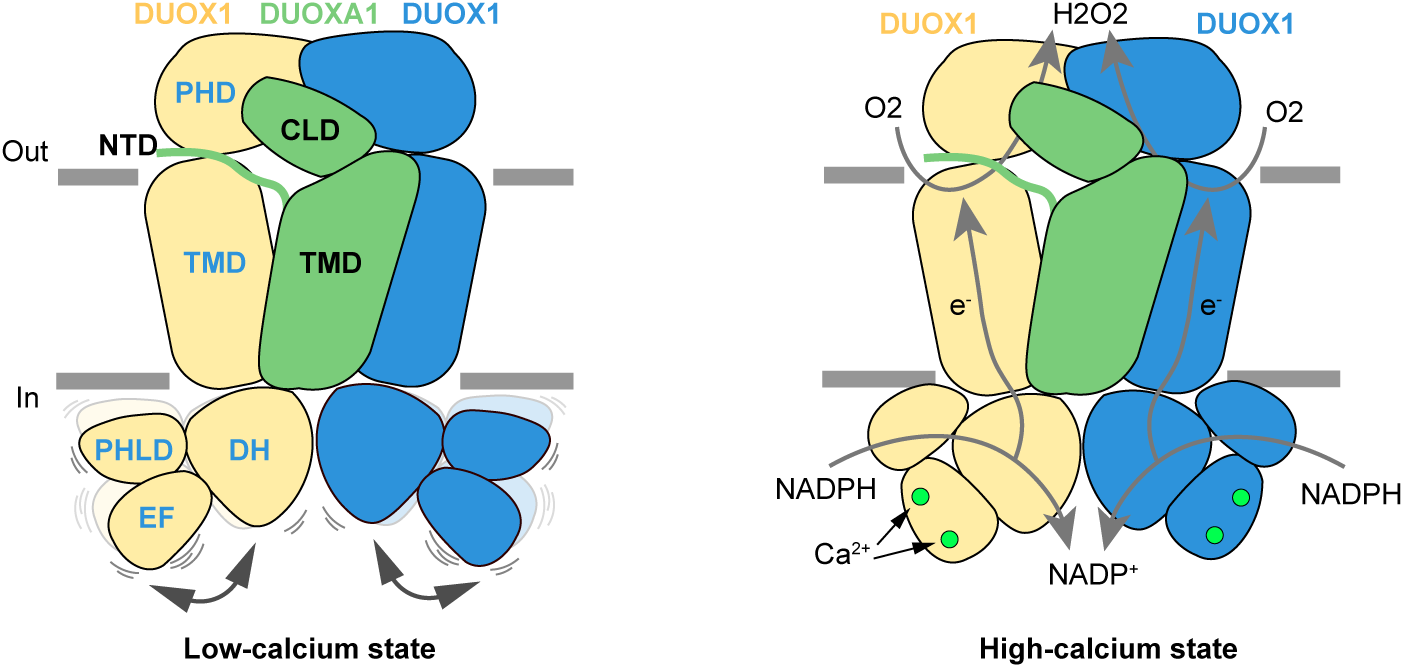
Activation mechanism of DUOX1 complex by calcium. Two DUOX1 and one DUOXA1 subunit are shown as cartoon and colored the same as Fig. 1d. Calcium ions are presented as green spheres. Electron transfer pathways are indicted with gray arrows.

## Materials and Methods

### Cell culture

HEK293F suspension cells (Thermo Fisher Scientific) were cultured in Freestyle 293 medium (Thermo Fisher Scientific) supplemented with 1% FBS at 37°C with 6% CO_2_ and 70% humidity. Sf9 insect cells (Thermo Fisher Scientific) were cultured in SIM SF (Sino Biological) at 27°C. The cell lines were routinely checked to be negative for mycoplasma contamination.

### Protein expression and purification

We constructed a modified BacMam vector ^25, 26^ with N-terminal GFP tag guided by rat FSHβ signal peptide ^9^ and cloned human DUOX1 into this vector. The cDNAs of human DUOXA1 were cloned into a non-tagged BacMam vector ^25, 26^. The two expression were further merged into one bicistronic vector by the LINK sequence on the modified vector ^25, 27^. The baculoviruses were produced using the Bac-to-Bac system and amplified in Sf9 cells. For protein expression, HEK293F cells cultured in Freestyle 293 medium at density of 2.8 × 10^6^ ml^-1^ were infected with 15% volume of P2 virus. 10 mM sodium butyrate was added to the culture 12 hours post infection and transferred to a 30°C incubator for another 36 hours before harvesting. Cells were collected by centrifugation at 4,000 rpm (JLA 8.1000, Beckman) for 10 min, and washed with 20 mM Tris (pH 8.0 at 4°C), 150 mM NaCl, 2 mM ethylene glycol tetraacetic acid (EGTA), 2 μg/ml aprotinin, 2 μg/ml pepstatin, 2 μg/ml leupeptin, flash frozen and storage at −80°C.

For each batch of protein purification, cell pellet corresponding to 0.5 liter culture was thawed and extracted with 20 ml Buffer A [20 mM Tris pH 8.0 at 4 °C, 150 mM NaCl, 5 μg/ml aprotinin, 5 μg/ml pepstatin, 5 μg/ml leupeptin, 20% (v/v) glycerol, 2 mM EGTA] containing 1 mM phenylmethanesulfonyl fluoride (PMSF) and 1% (w/v) digitonin (Biosynth) at 4°C for 50 min. 1 mM iodoacetamide (Sigma – I1149) was added during the detergent extraction procedure to reduce non-specific cysteine crosslinking. The supernatant was ultra-centrifuged at 50,000 rpm (TLA100.3, Beckman) for 50 min. The solubilized proteins were loaded onto 5 ml Streptactin Beads 4FF (Smart-Lifesciences) column and washed with 20 ml Buffer A+ 0.1% digitonin. The column was washed with 100 ml Buffer A + 0.1% digitonin plus 10 mM MgCl_2_ and 1 mM adenosine triphosphate (ATP) to remove contamination of heat shock proteins. Then the column was washed with 40 ml Buffer A + 0.1% digitonin again to remove residual MgCl_2_ and ATP. The target protein was assembled into the peptidisc on the Streptactin Beads through applying 4 ml 1 mg/ml NSPr in 20 mM Tris pH 8.0 ^11^. Then the column was washed with 100 ml Buffer A to remove free NSPr. The assembled peptidiscs were eluted with 40 ml Buffer A + 5 mM D-desthiobiotin (IBA). Eluted protein was concentrated using 100-kDa cut off concentrator (Millipore) and further purified by Superose 6 increase (GE Healthcare) running in HBS (20 mM Hepes pH 7.5, 150 mM NaCl) + 0.5 mM EGTA. Fraction 19 corresponding to DUOX1 + DUOXA1 peptidisc complex was concentrated to A280/415 = 4.4/1.4 with estimated concentration of 10.7 μM DUOX1 subunits (ε_415_=0.131 μM^−1^ cm^−1^).

### Enzymatic assay

The membrane fractions of DUOX1 for enzymatic assay were prepared as previously reported with minor modification ^28^. Briefly, cells were washed with Buffer A. After centrifuging at 4,000 rpm for 10 min at 4 °C, the cell pellets were broken using a needle for 12 times in 1 ml of 20 mM Tris pH 8.0 at 4 °C containing 0.1 mM dithiothreitol, 10 mM EGTA (pH 8.0) and the mixture of protease inhibitors. The pellet was removed by centrifuging at 7,000 rpm for 15 min, and the supernatant was collected and then centrifuged at 65,000 rpm (TLA100.3, Beckman) for 1 hour. The membrane pellet was resuspended in HBS containning 1 mM EGTA. Meanwhile, 10 μl membrane was solubilized by 100 μl TBS+1% digitonin with the mixture of protease inhibitors for 1 h at 4 °C for FSEC. The protein concentrations in the membrane were estimated by comparing their GFP fluorescence signal to that of a purified GFP-tagged DUOX1 complex.

The H_2_O_2_-generating activity of DUOX1 complex was determined using the amplex red assay ^29^. The concentrations of H_2_O_2_ solution were determined by measuring UV-Vis absorbance at 240 nm with spectrophotometer (Pultton) and calculated using molar extinction coefficient of 43.6 M^−1^ cm^−1^. The concentration of H_2_O_2_ solution was further validated by reacting with Amplex red to generate resorufin which has ε_571_=69,000 M^−1^ cm^−1^ ^29^. Then the H_2_O_2_ solution with known concentration was used to calibrate the resorufin fluorescence curve (excitation, 530 nm; emission, 590 nm) measured using a Microplate Reader (BioTek Synergy HT) at 37 °C.

The H_2_O_2_-generating reaction of the membrane fraction containing DUOX1 complex was performed at 37 °C in 0.15 ml of HBS with 1 mM EGTA, 10 μM FAD, 100 μM NADPH, 50 μM amplex red, 0.067 mg/ml horseradish peroxidase and 0.0576 mg/ml SOD. Ca^2+^ concentrations were determined using fluorescent indicators fura-2 or fluo3-FF. The K_cat_ and K_m_ values of the membrane fraction containing DUOX1 complex were determined at 37 °C with different concentrations of NADPH in the presence or absence of 1.4 mM CaCl_2_. The H_2_O_2_-generating reaction of the purified DUOX1 complex in peptidisc was performed at 27 °C in 0.15 ml of HBS + 1 mM EGTA, 10 μM FAD, 100 μM NADPH, 50 μM amplex red, 0.067 mg/ml horseradish peroxidase and 0.0576 mg/ml SOD in the presence or absence of 1.1 mM CaCl_2_. Progress of the reactions was monitored continuously by following the increase of the resorufin fluorescence, and the initial rates were calculated from the linear slopes of the progress curves. The activity of DUOX1 complex was determined by minusing the background of the corresponding buffer without DUOX1 samples.

### Cryo-EM sample preparation and data acquisition

The peptidisc sample was supplemented with 2.5 mM EGTA (low-calcium) or 0.5 mM free calcium (high-calcium) for cryo-EM analysis, respectively. Both samples contain 100 μM FAD as the cofactor and 500 μM NADPH as the substrate. To overcome the preferred orientation problem, 0.5 mM non-solubilizing detergent fluorinated octyl-maltoside was added to the sample before cyro-EM sample preparation. Aliquots of 1.5 μL protein sample were placed on graphene oxide (GO) coated grids as previously reported ^30^. Grids were blotted for 3 s at 100% humidity and flash-frozen in liquid ethane cooled by liquid nitrogen using Vitrobot Mark I (FEI). Grids were then transferred to a Titan Krios (FEI) electron microscope that was equipped with a Gatan GIF Quantum energy filter and operated at 300 kV accelerating voltage. Image stacks were recorded on a Gatan K2 Summit direct detector in super-resolution counting mode using SerialEM at a nominal magnification of 130,000 × (calibrated pixel size of 1.045 Å/pixel), with a defocus ranging from −1.5 to −2.0 μm. Each stack of 32 frames was exposed for 7.12 s, with a total dose about 50 e^-^/ Å ^2^ and a dose rate of 8 e^-^/pixel/s on detector.

### Image processing

The image processing workflow is illustrated in Fig. S2 and Fig. S4. A total of 7,076 super-resolution movie stacks of the high-calcium state sample and 2,076 stacks of the low-calcium state sample were collected using Serial EM, and motion-corrected, dose weighted and two-fold binned to a pixel size of 1.045 Å using MotionCor2 ^31^. Contrast transfer function (CTF) parameters were estimated with Gctf ^32^. Micrographs with ice or ethane contamination, and empty carbon were removed manually. Autopicking were performed using Gautomatch (kindly provided by Kai Zhang). All subsequent classification and reconstruction was performed in Relion 3.1 ^33^ unless otherwise stated. Reference-free 2D classification was performed to remove contaminants. Initial model was generated using cryoSPARC ^34^. Particles were subjected to multi-reference 3D classification ^35, 36^ and random-phase 3D classification ^35, 36^. Phase-randomized models were generated from the model obtained from previous refinement using randomize software (from the lab of Nikolaus Grigorieff). Further CTF refinement was then performed with Relion 3.1 using C2 symmetry. The particles were then re-extracted, re-centered, and re-boxed from 256 pixels to 320 pixels for consensus refinement. To improve the density of cytosolic layer, particles were symmetry expended ^12^ for multibody refinement^13^. One soft mask (the large body) that covers the extracellular domain together with transmembrane domain of one protomer was generated from the consensus map using UCSF Chimera and Relion 3.1^37^. The other soft mask (the small body) covers the cytosolic domains of the same protomer. 3D multi-body refinements ^13^ were performed using the two soft masks and the parameters determined from previous consensus refinement. The motions of the bodies were analyzed by relion_flex_analyse in Relion 3.1^37^. The two half-maps of each body generated by 3D multi-body refinement were subjected to post-processing in Relion 3.1^37^. The masked and sharpened maps of each body were aligned to the consensus map using UCSF Chimera^37^ and summed using Relion 3.1^37^ to generate the composite maps for visualization and model building. All of the resolution estimations were based on a Fourier shell correlation (FSC) of 0.143 cutoff after correction of the masking effect. B-factor used for map sharpening was automatically determined by the post-processing procedure in Relion 3.1 ^33^. The local resolution was estimated with Relion 3.1 ^33^.

### Model building

The composite maps derived from multibody refinement were used for model building. The structures of PHD, TMD, EF1-2 and DH domains of hDUOX1 were generated using phyre2 server ^38^ based on PDB ID: 6ERC, 5O0T, 4IL1 and 5O0X, and manually docked into the cryo-EM maps using Chimera ^37^. Initial models of PHLD were generated by Rosetta Web Server using *ab initio* mode ^39^, manually selected according to the distances calculated by RaptorX Contact Prediction server ^40^ and validated by the fitting between model and cryo-EM densities, especially the location of bulky aromatic residues. The partial model of hDUOXA1 were generated using EM builder ^41^. The initial models were iteratively built using Coot ^42^ and refined using Phenix in real space ^43^.

### Data availability

Data supporting the findings of this manuscript are available from the corresponding author upon reasonable request. The cryo-EM map of DUOX1-DUOXA1 in the high-calcium and low-calcium states have been deposited in the EMDB under ID code EMD-30556 and EMD-30555. The atomic coordinate of DUOX1-DUOXA1 in the high-calcium and low-calcium states have been deposited in the PDB under ID code 7D3F and 7D3E, respectively.

## Acknowledgement

We thank Yunlu Kang, Wenjun Guo, Yange Niu, Dian Ding, Mengmeng Wang, Miao Wei and Xiao He for illustration. We thank Miao Wei for making the rat FSHβ secretion signal-guided GFP vector. We thank Prof. Jiahuai Han for providing the cDNA of hDUOX1 and hDUOXA1, Prof. Helmut Grasberger for providing the cDNA of hDUOX2 and hDUOXA2, Prof. Yuji Kohara for providing the cDNA of c.elegans duox-2, bli-3 and doxA1. Cryo-EM data collection was supported by Electron microscopy laboratory and Cryo-EM platform of Peking University with the assistance of Xuemei Li, Daqi Yu, Xia Pei, Bo Shao, Guopeng Wang, and Zhenxi Guo. Part of structural computation was also performed on the Computing Platform of the Center for Life Science and High-performance Computing Platform of Peking University. This work is supported by grants from the Ministry of Science and Technology of China (National Key R&D Program of China, 2016YFA0502004 to L.C.), National Natural Science Foundation of China (91957201, 31622021, 31870833 and 31821091 to L.C., 31900859 to J.-X. W.), Beijing Natural Science Foundation (5192009 to L.C.), and Young Thousand Talents Program of China to L.C., and the China Postdoctoral Science Foundation (2016M600856, 2017T100014, 2019M650324, and 2019T120014 to J.-X.W.). J.-X. W. is supported by the Boya Postdoctoral Fellowship of Peking University and the postdoctoral foundation of the Peking-Tsinghua Center for Life Sciences, Peking University (CLS).

## Author contributions

Lei Chen initiated the project. Jing-Xiang Wu purified proteins and prepared the cryo-EM samples, collected the cryo-EM data and processed the cryo-EM data. Lei Chen built and refined the atomic model. Rui Liu and Kangcheng Song screened NOX constructs. All authors contributed to the manuscript preparation.

---

**Fig. S1.**
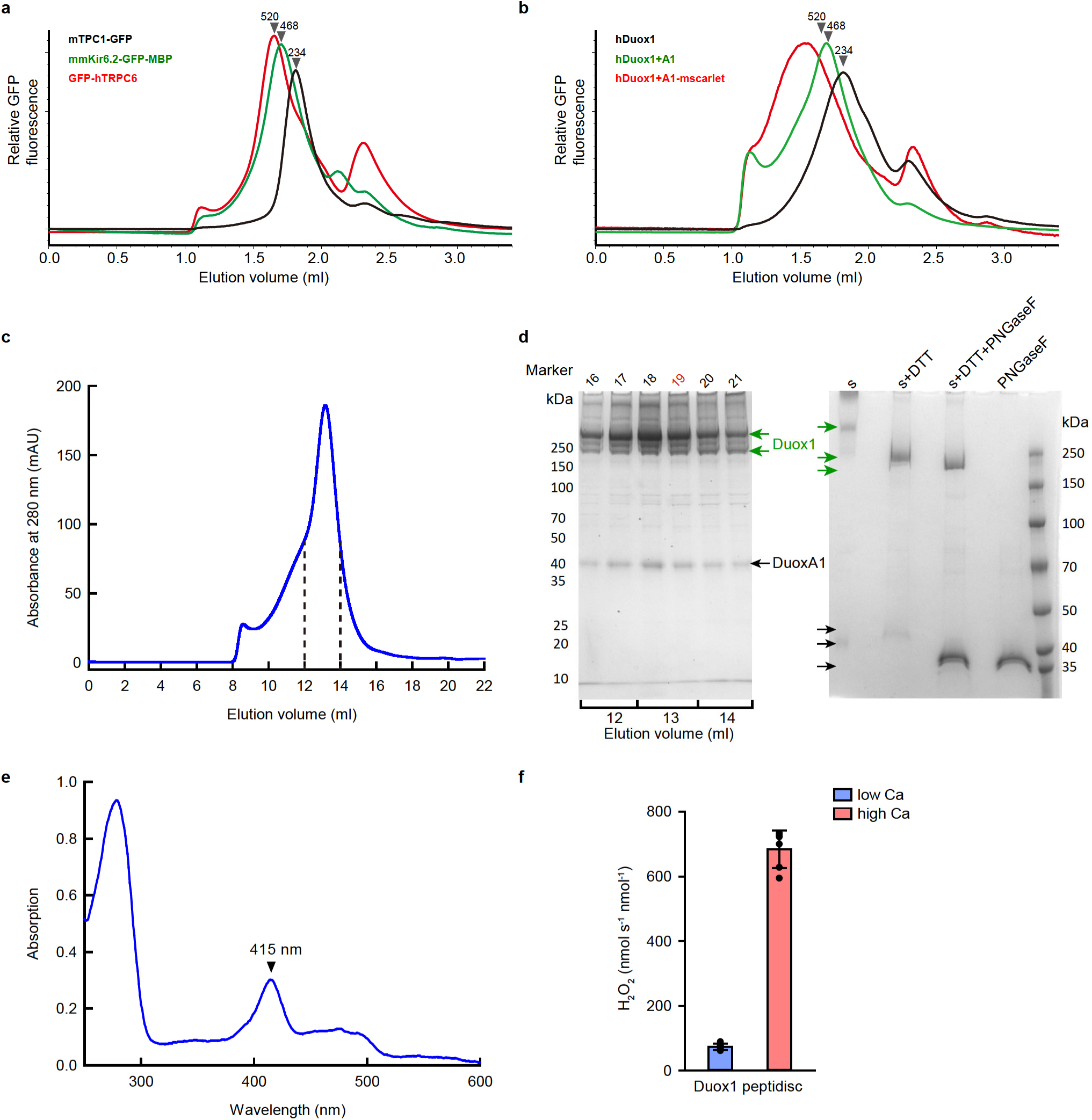
Biochemical characterization of hDUOX1-hDUOXA1 complex. **a**, FSEC traces of three membrane proteins as molecular weight markers. mTPC1-GFP (234 kDa): mouse TPC1 channel with C-terminal GFP; mmKir6.2-MBP-GFP (468 kDa): Mus musculus Kir6.2 channel with C-terminal MBP-GFP; GFP-hTRPC6 (520 kDa): human TRPC6 with N-terminal GFP. **b**, FSEC traces of hDUOX1 and hDUOX1-hDUOXA1 complex. The positions of the molecular weight markers described in **a** are indicated by arrowheads. hDUOX1: hDUOX1 with N-terminal GFP; hDUOX1+A1: hDUOX1 with N-terminal GFP was coexpressed with nontagged hDUOXA1; hDUOX1+A1-mscarlet: hDUOX1 with N-terminal GFP was coexpressed with hDUOXA1 with C-terminal MBP and mscarlet. **c**, Size-exclusion chromatography of hDUOX1-hDUOXA1 peptidisc complex on a superose 6 column. The fractions indicated by dashed lines were used for SDS–PAGE. **d**, SDS–PAGE of hDUOX1-hDUOXA1 protein samples. SDS–PAGE (left) of the size-exclusion chromatography fractions labeled in **a**. Fraction 19 highlighted in red was concentrated for cryo-EM sample preparation. SDS–PAGE (right) of hDUOX1-hDUOXA1 protein samples treated with DTT or DTT+PNGase F. The positions of hDUOX1 and hDUOXA1 subunits are indicated by green and black arrows, respectively. **e**, UV-Vis spectra of the purified hDUOX1-hDUOXA1 complex. The positions of the Soret peaks are indicated by arrowheads. **f**, Activity of the purified hDUOX1-hDUOXA1 complex peptidisc sample in the presence or absence of calcium. Mean ± s.d., n = 6 biologically independent samples.

**Fig.S2.**
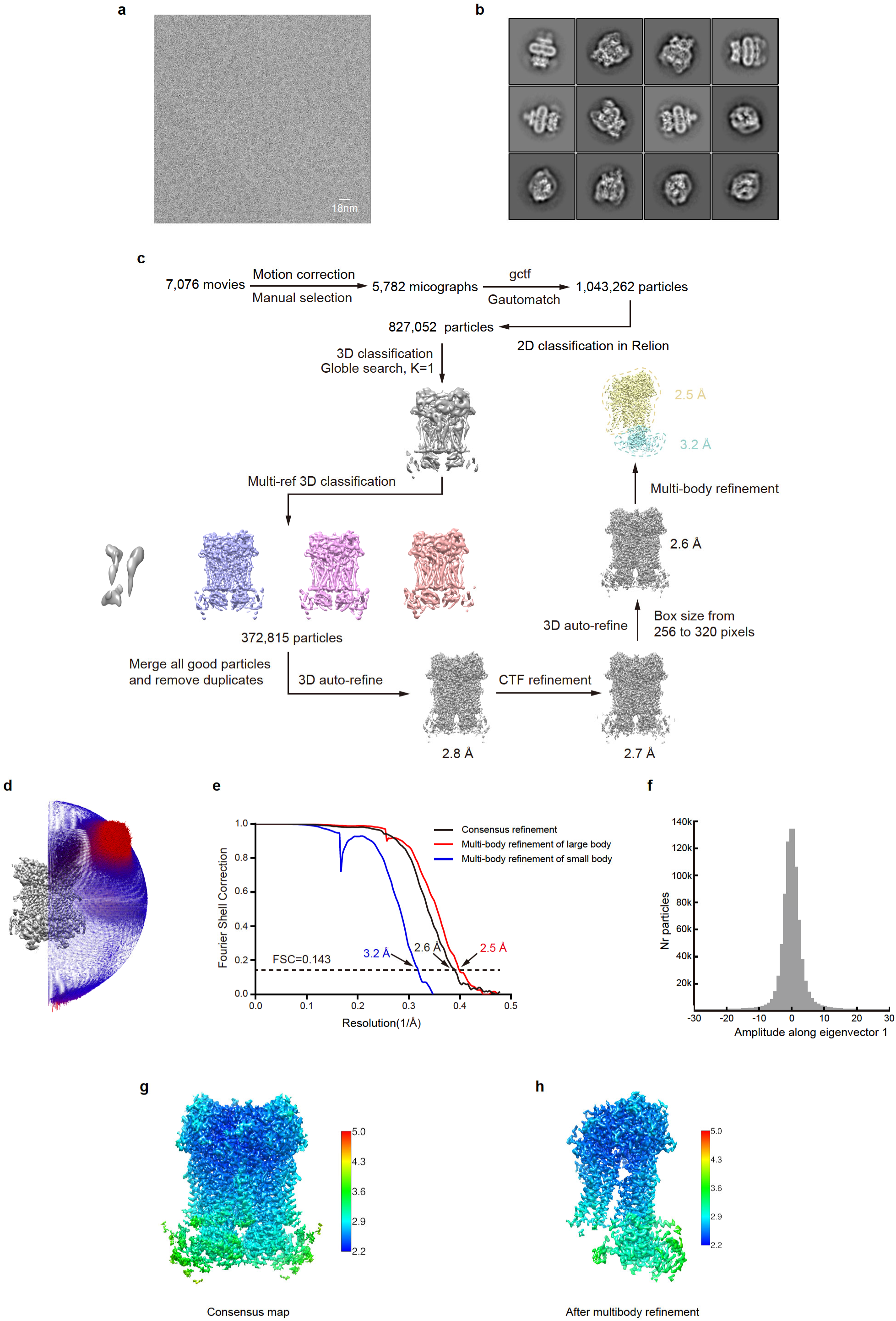
Cryo-EM image processing of hDUOX1-hDUOXA1 complex in the high-calcium state. **a**, Representative raw micrograph of hDUOX1-hDUOXA1 in the high-calcium state. **b**, Representative 2D class averages of hDUOX1-hDUOXA1 in the high-calcium state. **c**, The cryo-EM data processing workflow for hDUOX1-hDUOXA1 in the high-calcium state. **d**, The angular distribution for the consensus refinement of hDUOX1-hDUOXA1 in the high-calcium state. **e**, Gold-standard FSC curves of hDUOX1-hDUOXA1 in the high-calcium state. Resolution estimations are based on the criterion of FSC 0.143 cutoff. **f**, Histogram of the amplitudes along the top eigenvector shows monomodal distribution of hDUOX1-hDUOXA1 in the high-calcium state. **g**, Local resolution distribution of the consensus map of hDUOX1-hDUOXA1 in the high-calcium state. **h**, Local resolution distribution of the composite map of hDUOX1-hDUOXA1 in the high-calcium state after multibody refinement.

**Fig. S3.**
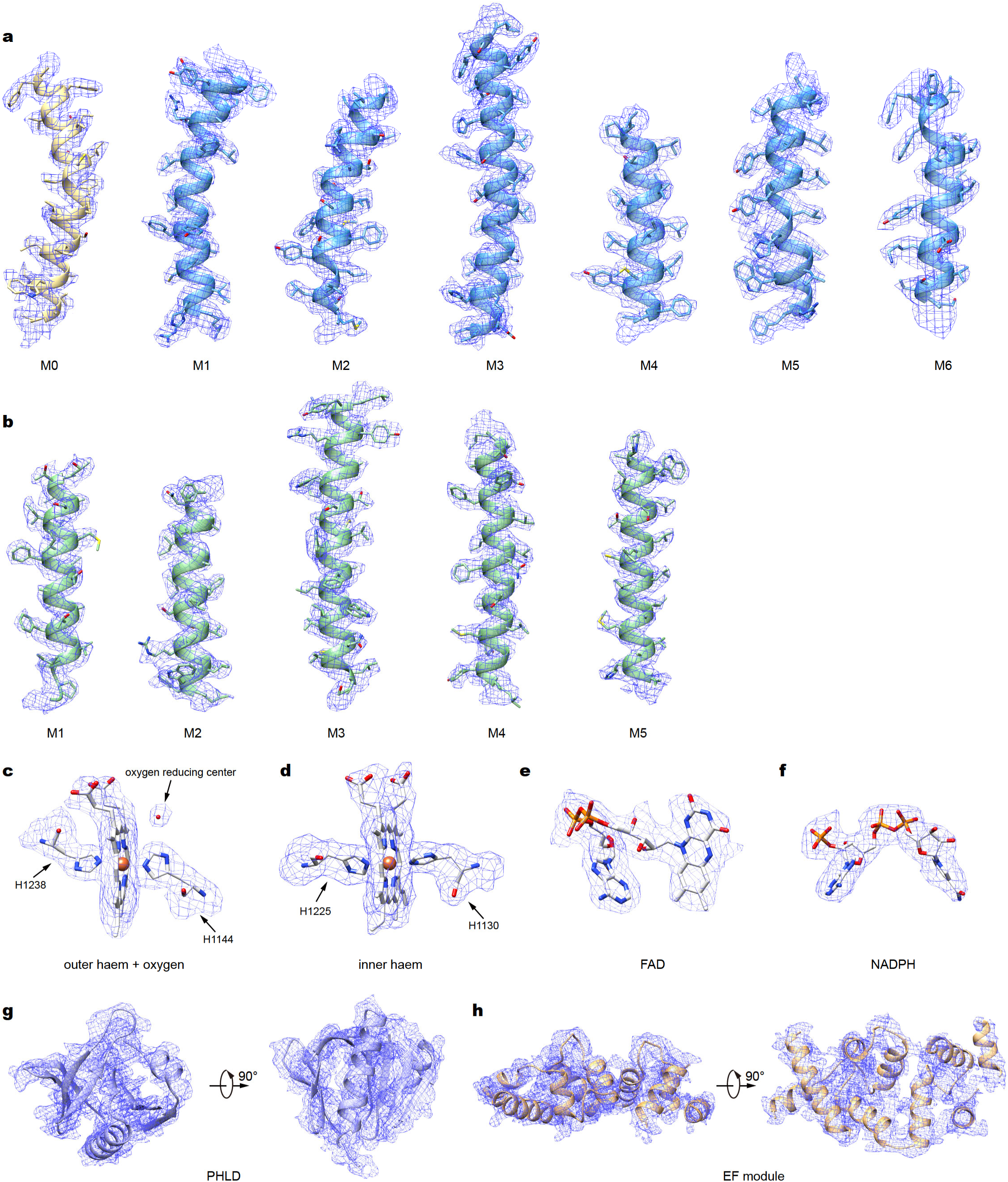
Representative electron densities of hDUOX1-hDUOXA1 complex in the high-calcium state. **a-b**, Representative densities of transmembrane helices of hDUOX1(**a**) and hDUOXA1(**b**). **c-f**, Representative densities of the outer haem with the interacted histidines and spherical density at the putative oxygen-reducing center (**c**), the inner haem with the interacted histidines (**d**), FAD (**e**) and NADPH (**f**). **g-h**, Representative densities of PHLD (**g**), EF module (**h**). Models of hDuox1 and hDuoxA1 are colored the same as in Fig. 1h. The representative densities are all shown in blue mesh. Electron density maps are all contoured at σ=1.86.

**Fig. S4.**
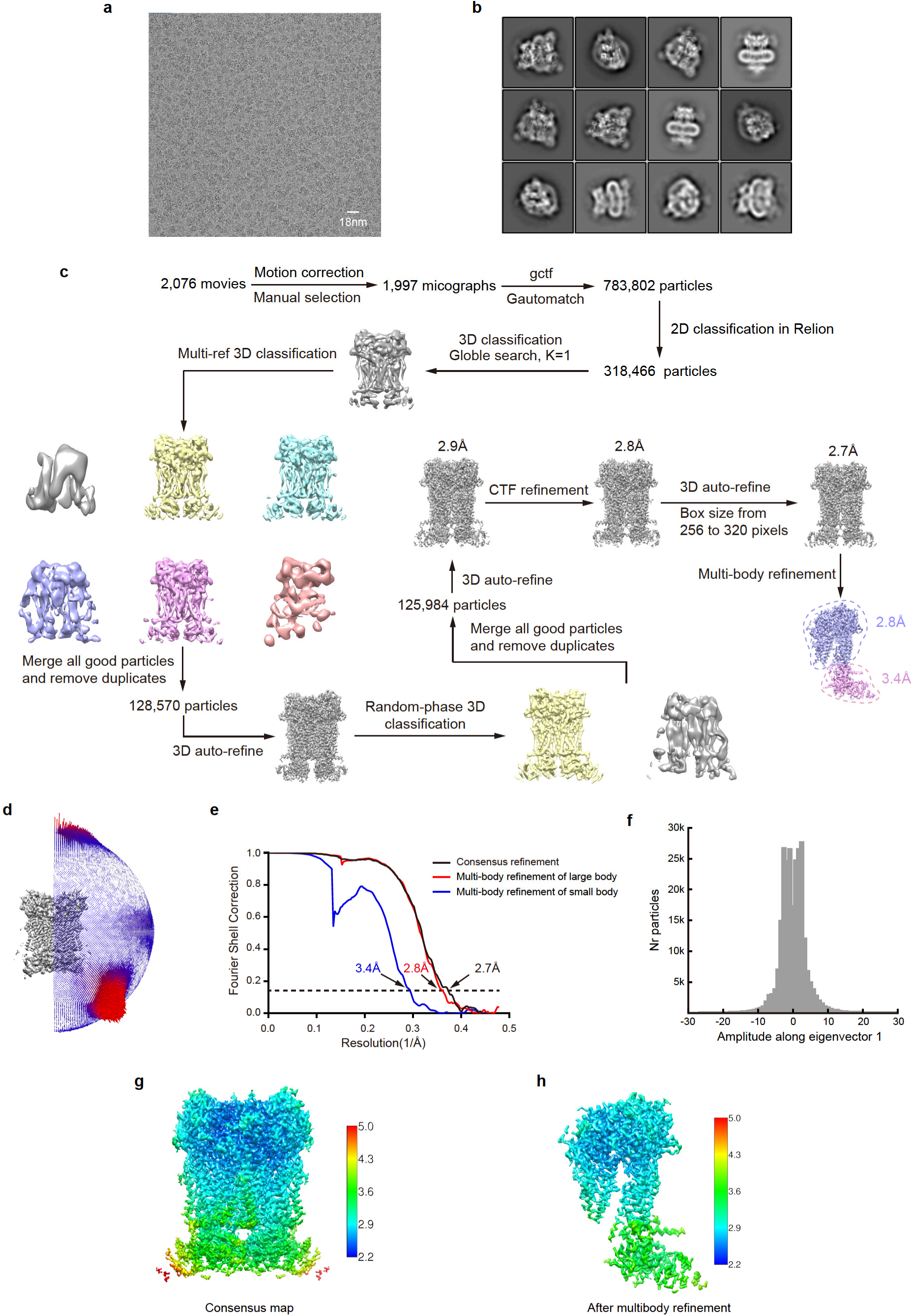
Cryo-EM image processing of hDUOX1-hDUOXA1 complex in the high-calcium state. **a**, Representative raw micrograph of hDUOX1-hDUOXA1 in the low-calcium state. **b**, Representative 2D class averages of hDUOX1-hDUOXA1 in the low-calcium state. **c**, The cryo-EM data processing workflow for hDUOX1-hDUOXA1 in the low-calcium state. **d**, The angular distribution for the consensus refinement of hDUOX1-hDUOXA1 in the low-calcium state. **e**, Gold-standard FSC curves of hDUOX1-hDUOXA1 in the low-calcium state. Resolution estimations are based on the criterion of FSC 0.143 cutoff. **f**, Histogram of the amplitudes along the top eigenvector of hDUOX1-hDUOXA1 in the low-calcium state shows a plateau-shaped distribution compared to that of the high-calcium state. **g**, Local resolution distribution of the consensus map of hDUOX1-hDUOXA1 in the low - calcium state. **h**, Local resolution distribution of the composite map of hDUOX1-hDUOXA1 in the low - calcium state after multibody refinement.

**Fig. S5.**
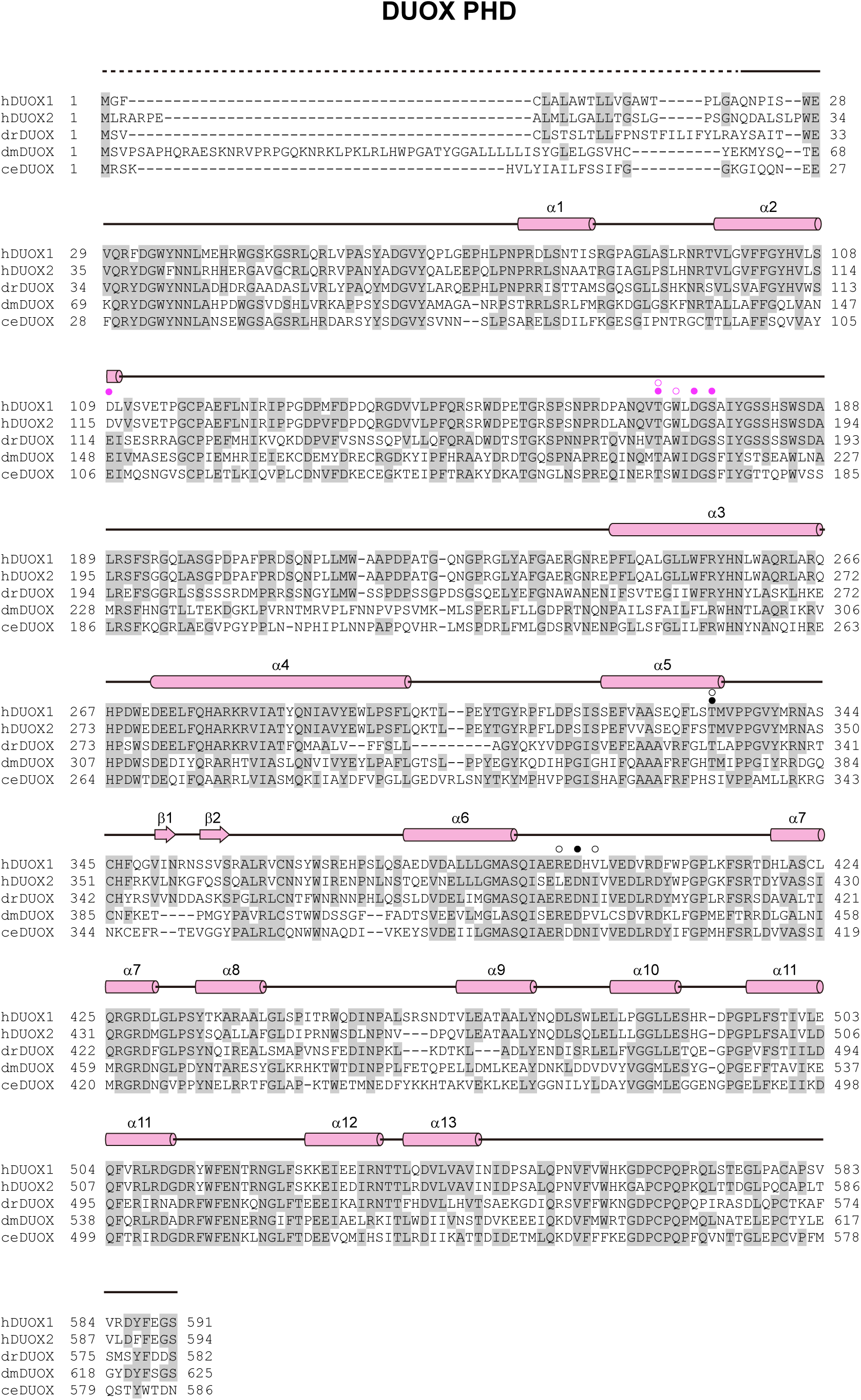
Sequence alignment of the peroxidase homology domain (PHD) of DUOX. The sequences of the *Homo sapiens* DUOX1 (hDUOX1), *Homo sapiens* DUOX2 (hDUOX2), *Danio rerio* DUOX (drDUOX), *Drosophila melanogaster* DUOX (dmDUOX) and *Caenorhabditis elegans* DUOX (ceDUOX) were aligned. Residues following PHD are omitted for clarity. The sequence alignment of Figs. S5-8 are all shown as follows: Conserved residues are highlighted in gray; Secondary structures are indicated as cylinders (α helices), arrows (β sheets) and lines (loops); Unmodeled residues are indicated as dashed line; The color of arrows and cylinders are the same as in Fig. 1h. Residues of cation binding site 1 (CBS1) and cation binding site 2 (CBS2) are indicated as black circles and violet circles, respectively. The filled circles and empty circles indicate side chain and main chain interactions.

**Fig. S6.**
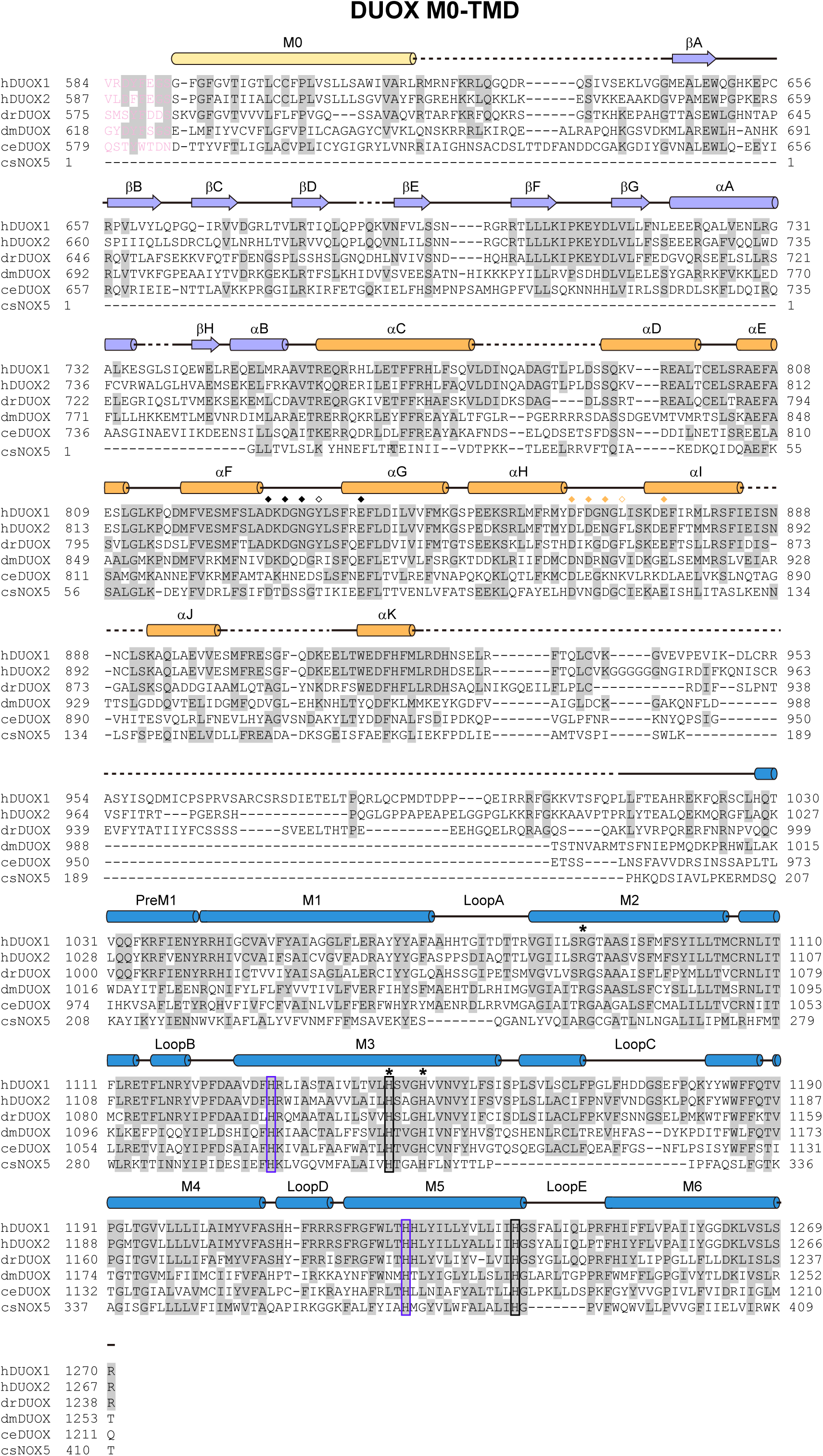
Sequence alignment from M0 helix to transmembrane domain (TMD) of DUOX. The sequences of the *Homo sapiens* DUOX1 (hDUOX1), *Homo sapiens* DUOX2 (hDUOX2), *Danio rerio* DUOX (drDUOX), *Drosophila melanogaster* DUOX (dmDUOX), *Caenorhabditis elegans* DUOX (ceDUOX) and *Cylindrospermum stagnale* NOX5 (csNOX5) were aligned. Residues belonging to PHD are shown in violet, and residues following TMD are omitted for clarity. Residues predicted to be responsible for calcium binding in EF1 and EF2 are indicated as black diamonds and orange diamonds, respectively. The filled diamonds and empty diamonds indicate side chain and main chain interactions. Histidines coordinating the outer haem and the inner haem are enclosed by black boxes and blue boxes, respectively. Residues of proposed oxygen substrate binding site are indicated as asterisks.

**Fig. S7.**
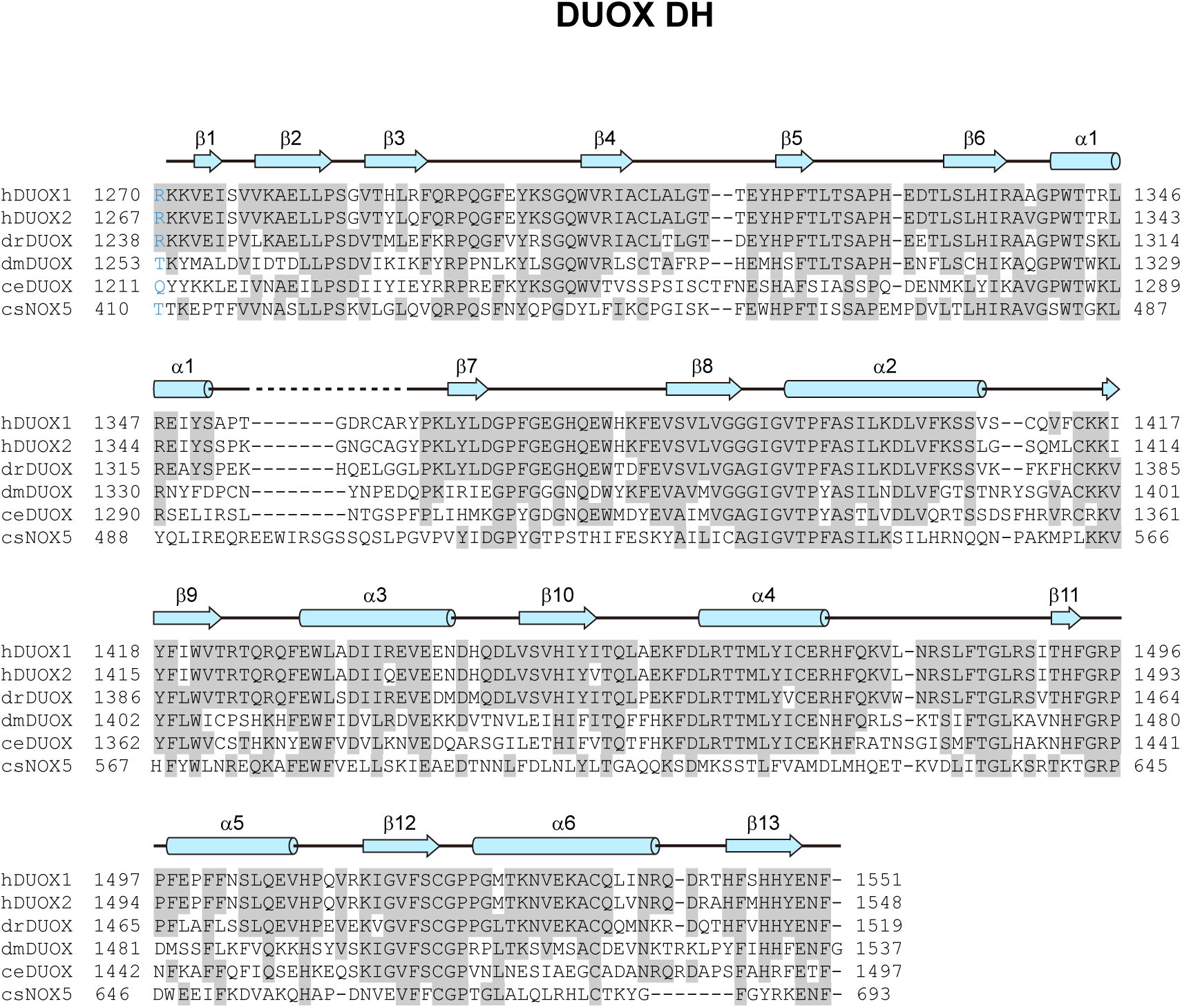
Sequence alignment of the dehydrogenase domain (DH) of DUOX. The sequences of the *Homo sapiens* DUOX1 (hDUOX1), *Homo sapiens* DUOX2 (hDUOX2), *Danio rerio* DUOX (drDUOX), *Drosophila melanogaster* DUOX (dmDUOX), *Caenorhabditis elegans* DUOX (ceDUOX) and *Cylindrospermum stagnale* NOX5 (csNOX5) were aligned. Residues belonging to TMD are shown in marine. Conserved residues are highlighted in gray. Secondary structures are indicated as cylinders (α helices), arrows (β sheets) and lines (loops). Unmodeled residues are indicated as dashed line. The color of arrows and cylinders are the same as in Fig. 1.

**Fig. S8.**
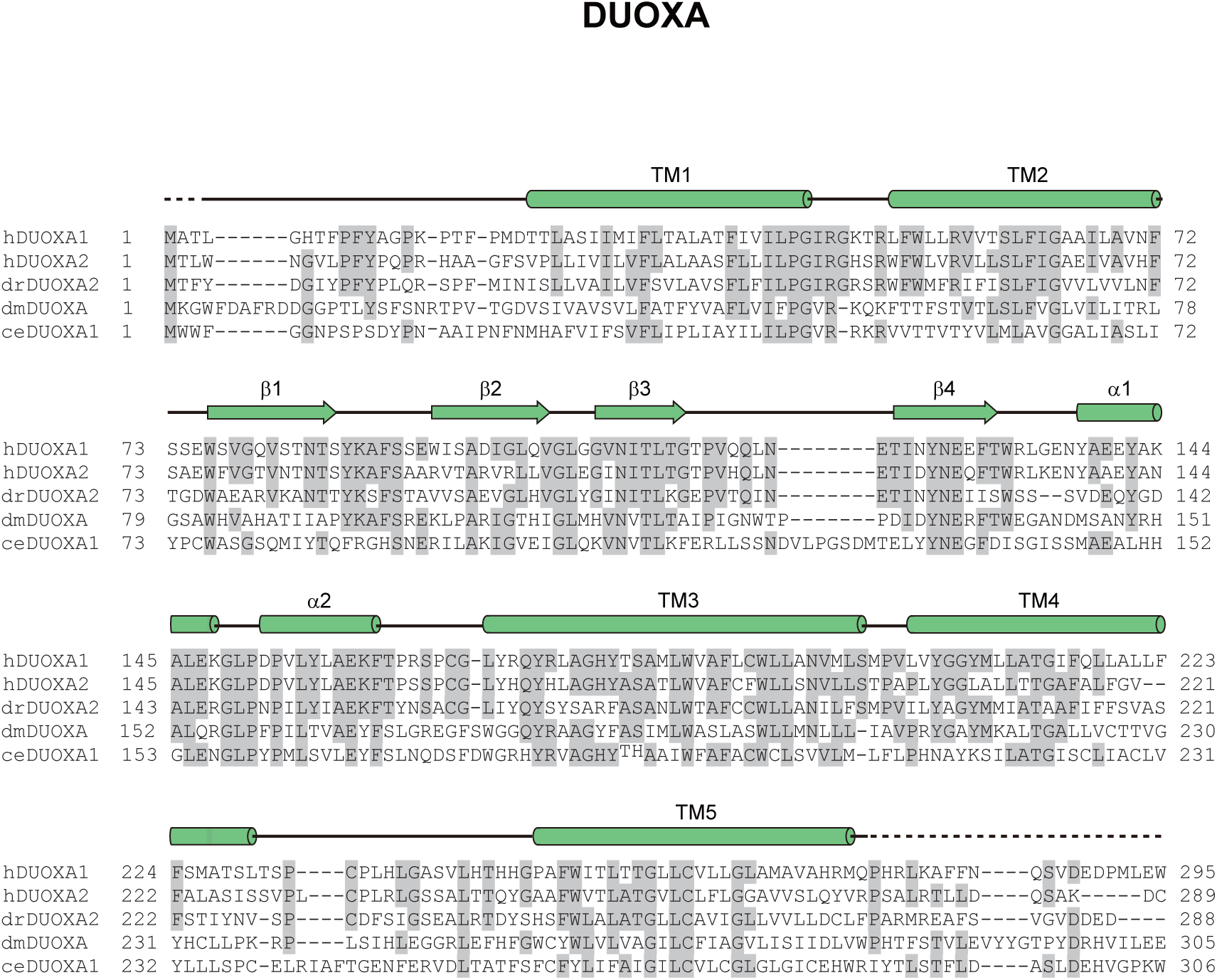
Sequence alignment of DUOXA. The sequences of the *Homo sapiens* DUOXA1. The sequences of the *Homo sapiens* DUOXA1 (hDUOXA1), *Homo sapiens* DUOXA2 (hDUOXA2), *Danio rerio* DUOXA2 (drDUOXA2), *Drosophila melanogaster* DUOXA (dmDUOXA) and *Caenorhabditis elegans* DUOXA1 (ceDUOXA1) were aligned. Conserved residues are highlighted in gray. C-terminal residues are omitted for clarity. Secondary structures are indicated as cylinders (α helices), arrows (β sheets) and lines (loops). Unmodeled residues are indicated as dashed lines. The color of arrows and cylinders are the same as in Fig. 1.

**Fig. S9.**
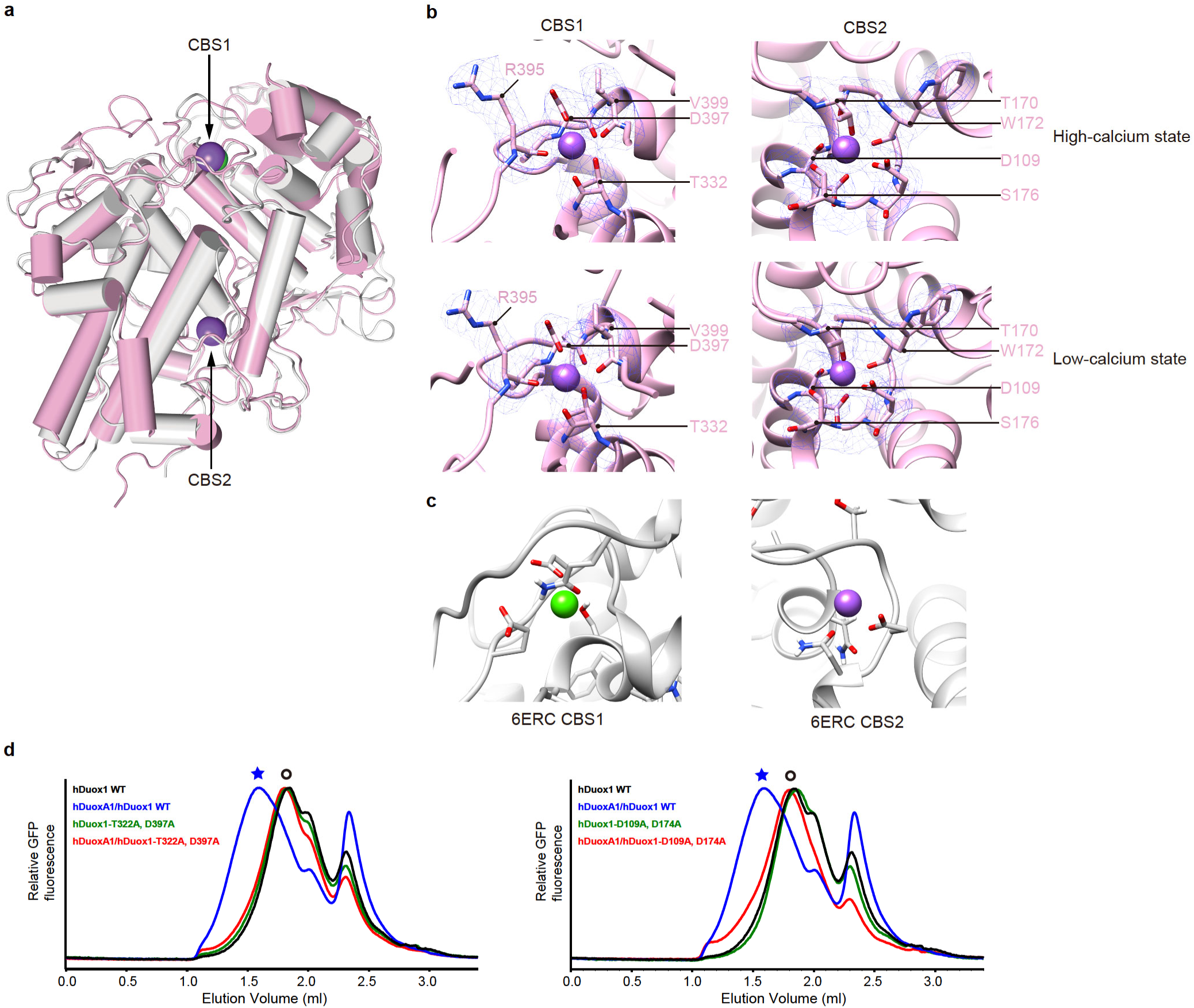
Structure of hDUOX1 PHD domain. **a,** Structural comparison between hDUOX1 PHD (pink) and DdPoxA (PDB ID: 6ERC, gray). CBS1 and CBS2 are denoted by arrows. **b,** Electron densities of CBS1 and CBS2 in the high-calcium state and low-calcium states. **c,** CBS1 and CBS2 in DdPoxA (PDB ID: 6ERC). **d,** FSEC profiles showed mutations of hDUOX1 CBS1 (T322A, D379A) or CBS2 (D109A, D174A) affect hDUOX1-hDUOXA1 tetramer assembly. Asterisks denote the peak position of tetrameric complex and circles denote hDUOX1 monomer peak position.

**Fig. S10.**
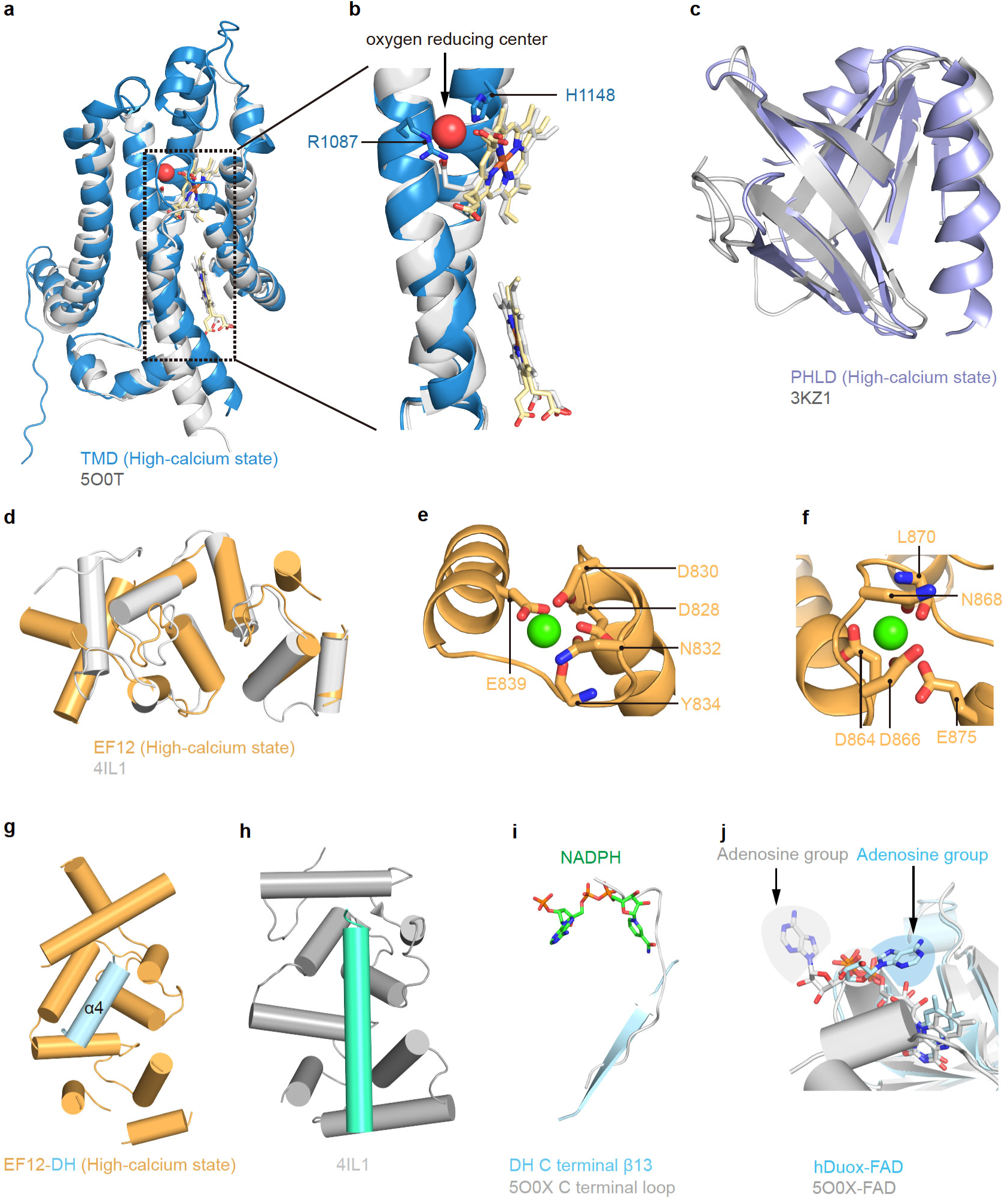
Structure of hDUOX1 TMD and cytosolic domains. **a,** Structural comparison between hDUOX1 TMD (blue) and csNOX (PDB ID: 5O0T, gray). The putative oxygen-reducing center is shown as red sphere. **b**, The close-up view of the similar architecture of the putative oxygen-reducing center. **c**, Structural comparison between hDUOX1 PHLD (light blue) and PDZ-RhoGEF (PDB ID:3KZ1, gray). **d**, Structural comparison between hDUOX1 EF module (orange) and Calcineurin B subunit (PDB ID: 4IL1, gray). **e**, Calcium binding on EF1 domain. Calcium ion is modeled according to 4IL1 and shown as green sphere. **f**, Calcium binding on EF2 domain. Calcium ion is modeled according to 4IL1 and shown as green sphere. **g**, Structure of hDUOX1 EF module (orange) in complex with α4 helix (light blue) of DH domain. **h**, Structure of calcineurin B subunit in complex with helix of calcineurin A subunit (PDB ID: 4IL1). **i**, Structural comparison of NADPH binding pocket between hDUOX1 (colored) and csNOX (PDB ID: 5O0X, gray). Only β13 and NADPH are shown for clarity. **j**, Structural comparison of FAD binding pocket between hDUOX1 (colored) and csNOX (PDB ID: 5O0X, gray). The large rotations of adenosine group of FAD are denoted by arrows.

**Fig. S11.**
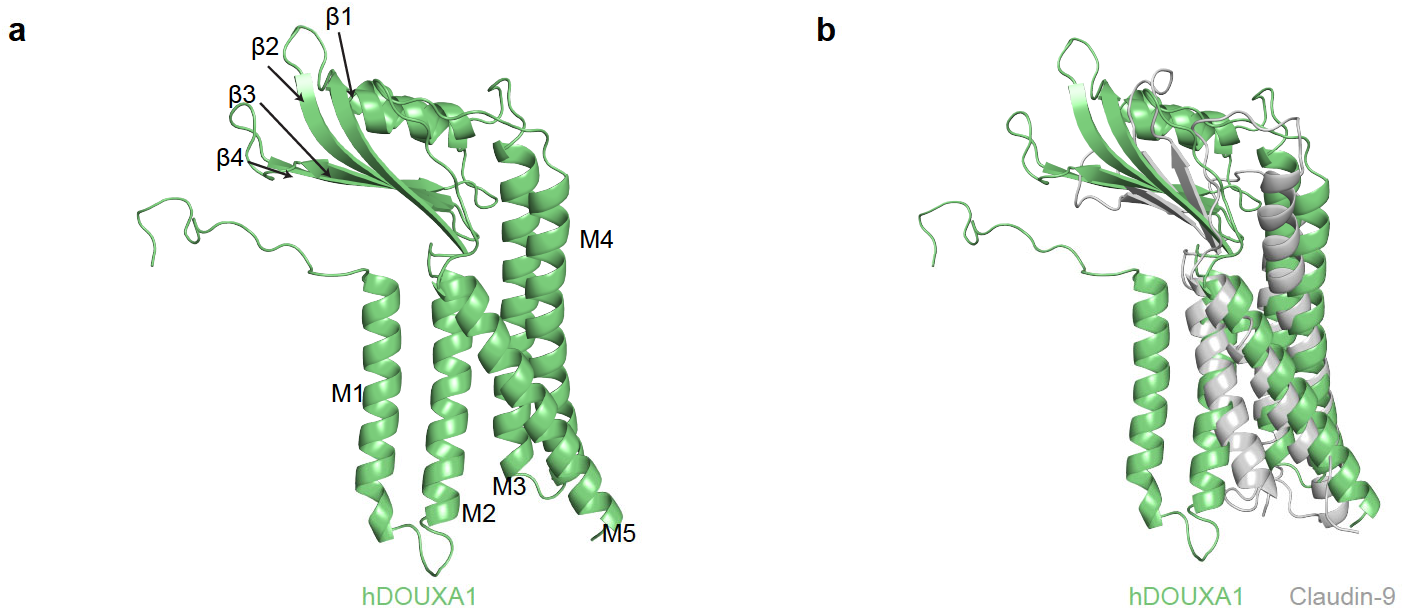
Structure of hDUOXA1 subunit. **a,** Structure of hDUOXA1 subunit with secondary structure labeled. **b**, Structural comparison of hDUOXA1 with claudin-9 (PDB ID:6OV2). hDUOXA1 is colored in green and claudin-9 is colored in gray, respectively.

**Movie S1 Structural changes of the cytosolic domains of hDUOX1-hDUOXA1 complex during calcium activation**

hDUOUX1-hDUOXA1 complex is shown as cartoon and colored the same as Fig.1h. Calcium ions are shown as green spheres. The movie starts from the whole molecule and then focuses on the cytosolic domains of one protomer. The structural changes are presented as a morph between the high-calcium state and the low-calcium state.

**Table. S1.**
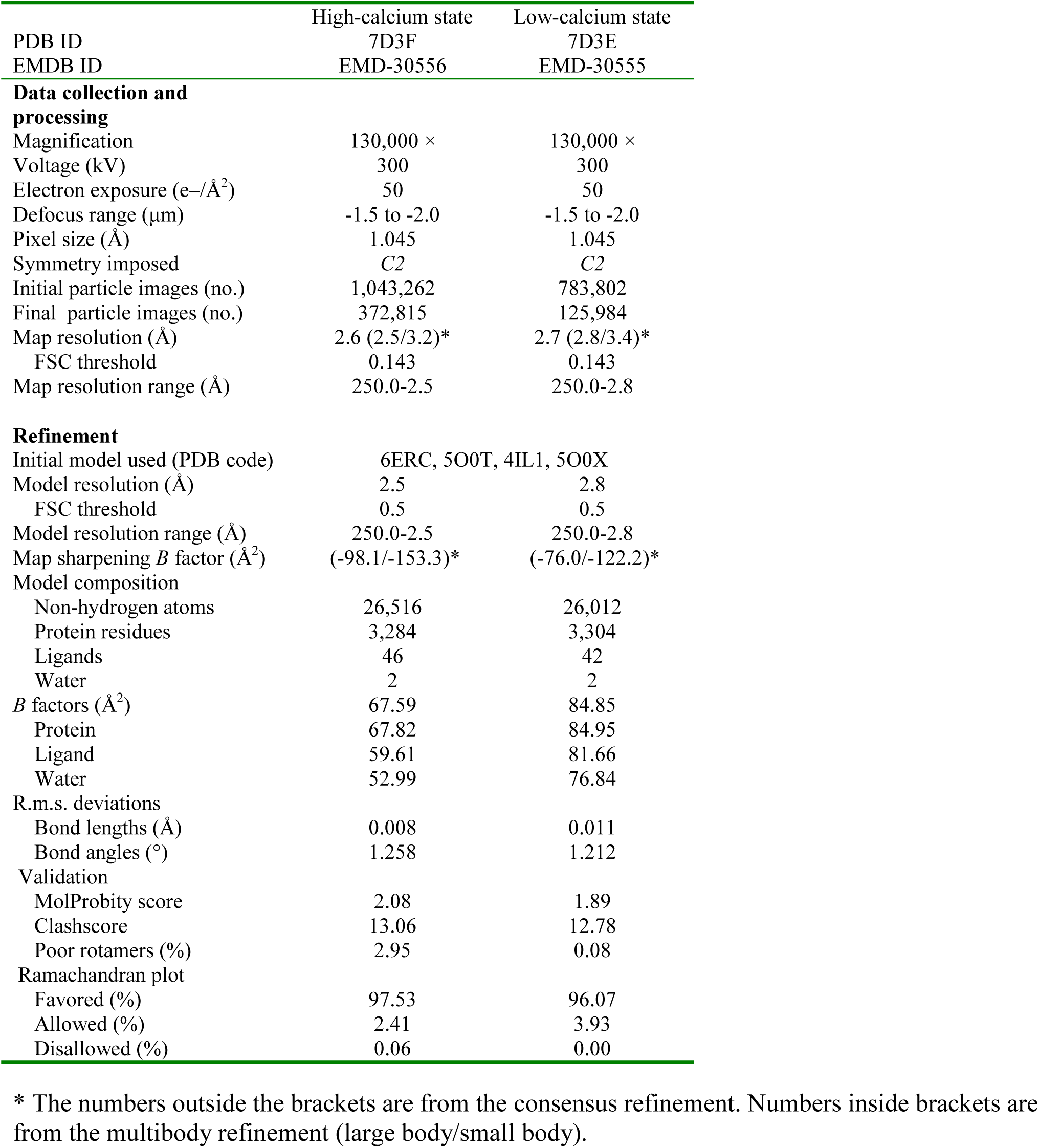
Cryo-EM data collection, refinement and validation statistics.

|  | High-calcium state | Low-calcium state |
| --- | --- | --- |
| PDB ID | 7D3F | 7D3E |
| EMDB ID | EMD-30556 | EMD-30555 |
| **Data collection and processing** |  |  |
| Magnification | 130,000 × | 130,000 × |
| Voltage (kV) | 300 | 300 |
| Electron exposure (e-/Å<sup>2</sup>) | 50 | 50 |
| Defocus range (μm) | -1.5 to -2.0 | -1.5 to -2.0 |
| Pixel size (Å) | 1.045 | 1.045 |
| Symmetry imposed | C2 | C2 |
| Initial particle images (no.) | 1,043,262 | 783,802 |
| Final particle images (no.) | 372,815 | 125,984 |
| Map resolution (Å) | 2.6 (2.5/3.2)* | 2.7 (2.8/3.4)* |
| FSC threshold | 0.143 | 0.143 |
| Map resolution range (Å) | 250.0-2.5 | 250.0-2.8 |
| **Refinement** |  |  |
| Initial model used (PDB code) | 6ERC, 5O0T, 4IL1, 5O0X |  |
| Model resolution (Å) | 2.5 | 2.8 |
| FSC threshold | 0.5 | 0.5 |
| Model resolution range (Å) | 250.0-2.5 | 250.0-2.8 |
| Map sharpening *B* factor (Å<sup>2</sup>) | (-98.1/-153.3)* | (-76.0/-122.2)* |
| Model composition |  |  |
| Non-hydrogen atoms | 26,516 | 26,012 |
| Protein residues | 3,284 | 3,304 |
| Ligands | 46 | 42 |
| Water | 2 | 2 |
| *B* factors (Å<sup>2</sup>) | 67.59 | 84.85 |
| Protein | 67.82 | 84.95 |
| Ligand | 59.61 | 81.66 |
| Water | 52.99 | 76.84 |
| R.m.s. deviations |  |  |
| Bond lengths (Å) | 0.008 | 0.011 |
| Bond angles (°) | 1.258 | 1.212 |
| Validation |  |  |
| MolProbity score | 2.08 | 1.89 |
| Clashscore | 13.06 | 12.78 |
| Poor rotamers (%) | 2.95 | 0.08 |
| Ramachandran plot |  |  |
| Favored (%) | 97.53 | 96.07 |
| Allowed (%) | 2.41 | 3.93 |
| Disallowed (%) | 0.06 | 0.00 |

\* The numbers outside the brackets are from the consensus refinement. Numbers inside brackets are from the multibody refinement (large body/small body).

